# Liver Receptor Homolog-1 (LRH-1/NR5A2) orchestrates hepatic inflammation and TNF-induced cell death

**DOI:** 10.1101/2023.05.24.542039

**Authors:** Rebekka Lambrecht, M. Eugenia Delgado, Vincent Gloe, Karina Schuetz, Anna Pia Plazzo, Barbara Franke, Truong San Phan, Jennifer Fleming, Olga Mayans, Thomas Brunner

## Abstract

Liver Receptor Homolog-1 (LRH-1/NR5A2) is a nuclear receptor that has been shown to promote apoptosis resistance in various tissues and disease contexts, however, its role in liver cell death remains unexplored. Deletion of LRH-1 in hepatocytes developed into a mild steatosis and inflammation already under steady-state conditions. Unexpectedly, hepatocyte-specific deletion of LRH-1 also resulted in a profound protection of mice from TNF-induced hepatocyte apoptosis and associated hepatitis. LRH-1-deficient hepatocytes showed elevated NF-ⲕB activity, while LRH-1 overexpression inhibited NF-ⲕB activity. This inhibition was based on direct physical interaction of the ligand-binding domain of LRH-1 and the Rel homology domain of NF-ⲕB subunit RelA. Mechanistically, we found that increased transcription of anti-apoptotic NF-ⲕB target genes, together with proteasomal degradation of pro-apoptotic BIM via regeneration-driven EGF receptor signaling, prevented mitochondrial apoptosis, ultimately protecting mice from TNF-induced liver damage. Collectively, our study demonstrates that LRH-1 is a critical modulator of cell death and inflammation in the healthy and diseased liver.

**Highlights:**

1. Hepatic LRH-1 deletion causes mild liver steatosis, fibrosis, and inflammation.
2. Female LRH-1-deficient mice are protected from TNF-induced liver damage.
3. LRH-1 interacts with NF-ⲕB and inhibits its activity.
4. LRH-1 deletion-provoked inflammation causes degradation of pro-apoptotic protein BIM.

**Graphical abstract:** 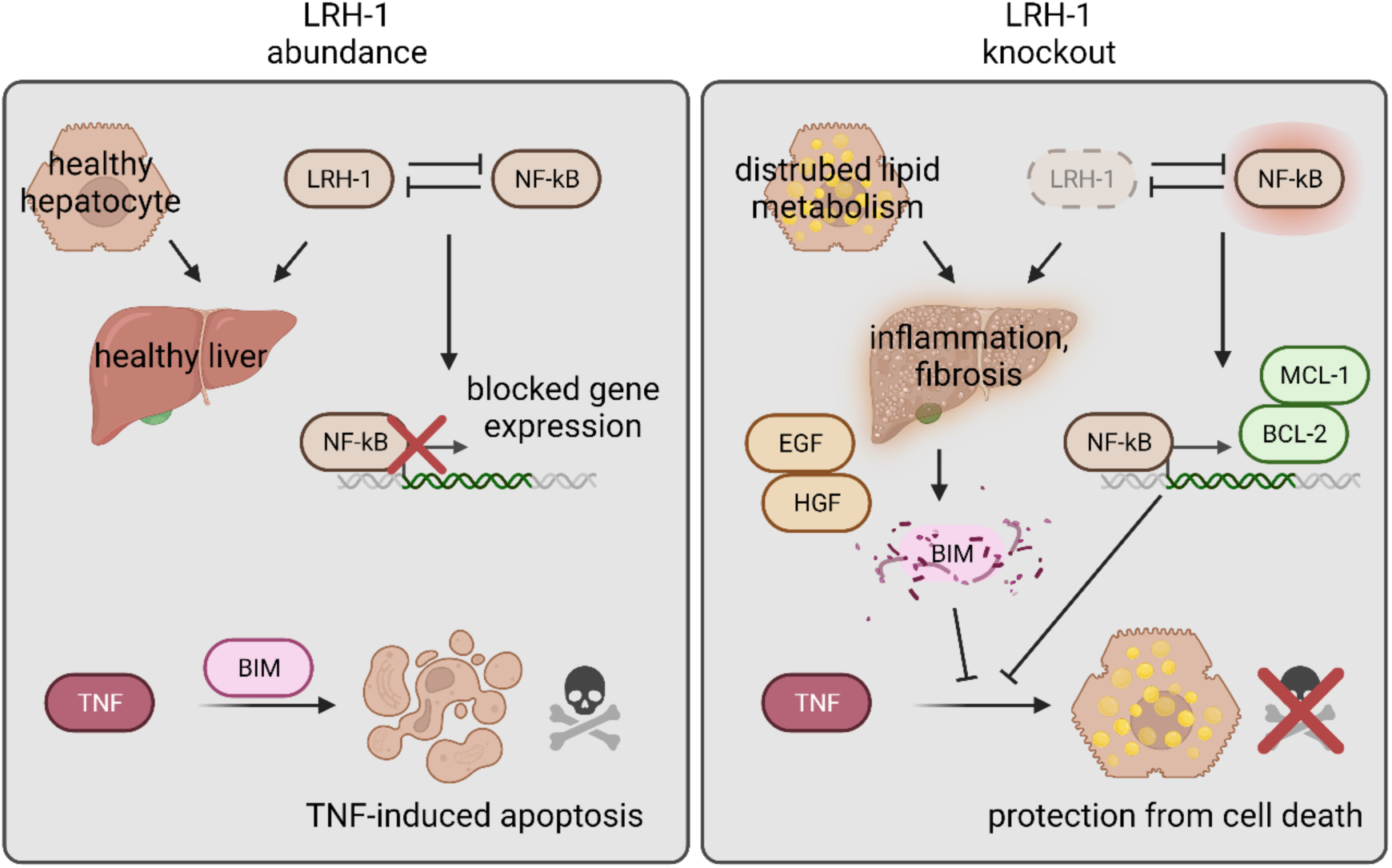

## Introduction

The nuclear receptor liver receptor homolog-1 (LRH-1/NR5A2) is highly expressed in tissues of endodermal origin and exhibits wide-ranging functions (Fernandez-Marcos et al., 2011; Sun et al., 2021). LRH-1 is essential during early development and differentiation, which is well documented by the embryonic lethality of mice with systemic LRH-1 deficiency (Paré et al., 2004). Furthermore, LRH-1 is a central regulator of proliferation in adult cells as it regulates the expression of cell cycle genes and co-activates other proliferation-regulating transcription factors (Botrugno et al., 2004; Schoonjans et al., 2005; Seitz et al., 2019; Wang et al., 2005). Notably, these proliferative functions, together with its high expression in several types of cancer, define LRH-1 as pro-tumorigenic factor (Nadolny and Dong, 2015).

Furthermore, LRH-1 is an established key regulator of various metabolic processes. LRH-1 controls the expression of enzymes involved in glycolysis, and amino acid and fatty acid metabolism, thereby sensing and regulating blood glucose levels and lipid and hormone abundancies (Hattori et al., 2014; Lu et al., 2000; Matsukuma et al., 2007; Miranda et al., 2018; Oosterveer and Schoonjans, 2014; Oosterveer et al., 2012; Xu et al., 2016). These processes are particularly important in the liver (summarized in Sun et al., 2021), albeit complex as for example both, mice with an LRH-1 gain-of-function mutation as well as mice with acute hepatic LRH-1 deletion develop hepatic steatosis (Miranda et al., 2018; Stein et al., 2017).

LRH-1 also critically modulates several inflammatory processes in different organs. Current research suggests that the role of LRH-1 in inflammation is highly tissue- and disease-specific, although in most conditions LRH-1 exerts anti-inflammatory and anti-apoptotic actions. LRH-1 counteracts pro-inflammatory cytokine signaling of the acute phase response in the liver by inducing the expression of IL-1 receptor antagonists (Venteclef et al., 2006). In the intestine, LRH-1 controls the local synthesis of anti-inflammatory glucocorticoids (Coste et al., 2007; Mueller et al., 2006) and supports the differentiation of anti-inflammatory macrophages (Lefèvre et al., 2015). Consequently, local LRH-1 deletion drastically exacerbates inflammatory bowel diseases (Coste et al., 2007; Mueller et al., 2006), while LRH-1 overexpression attenuates colitis in mice and cell death in patient-derived intestinal organoids (Bayrer et al., 2018). Treatment with an LRH-1 agonist prevented also pancreatic inflammation and protected β-islets cells from apoptosis (Cobo-Vuilleumier et al., 2018a; Martin Vázquez et al., 2022). The anti-apoptotic potential of LRH-1 was recently reinforced by showing that pharmacological inhibition of LRH-1 also promoted cell death in leukemic cells (Michalek et al., 2022). In contrast, LRH-1 appears to regulate the expression of pro-inflammatory cytokines in T cells and macrophages, and its pharmacological inhibition results in decreased immune cell-mediated liver damage (Schwaderer et al., 2017, 2020; Seitz et al., 2019).

Despite extensive characterization of LRH-1-regulated processes in development, proliferation, metabolism, and inflammation, only little is known about the role of LRH-1 in hepatocyte cell death. The liver is a central metabolic and storage organ. The direct connection with the intestine via the portal vein exposes the liver to potentially harmful substances and inflammatory mediators. Thus, the hepatic environment is required to maintain a tight balance between pathogen clearance during infections and inflammatory responses, which may cause hepatocyte cell death and loss of liver function. Here, we sought to elucidate the role of LRH-1 in inflammation-driven hepatic cell death by analyzing different experimental acute hepatitis models in mice with liver-specific LRH-1 deletion. Surprisingly, we found a paradoxical role of LRH-1 in promoting TNF-induced mitochondrial apoptosis via the regulation of the BCL-2 family.

## Results

### Hepatic LRH-1 deficiency provokes steady-state steatosis and inflammation

In the present study, we used mice with hepatocyte-specific deletion of LRH-1 (LRH-1 AlbCre, ‘LRH-1-deficient’) and their littermates with floxed LRH-1 gene, but without Cre expression, as control mice (LRH-1 L2L2) (Mataki et al., 2007). When we characterized the livers of untreated LRH-1 AlbCre mice, we found that in particular female LRH-1-deficient mice show pronounced lipid accumulation in the liver tissue and changed lipid distribution in isolated hepatocytes (Figure 1A, S1A). It was previously described that only acute deletion of LRH-1 and not the constitutive deletion results in hepatic steatosis, however, only male mice were used in their study (Miranda et al., 2018). We observed furthermore that LRH-1-deficient livers of both sexes show significantly increased collagen deposition (Figure 1B) and elevated serum alanine aminotransferase (ALT) values (Figure 1C), indicating ongoing hepatocyte cell death and compensatory scar tissue formation. To better understand this pathological condition in LRH-1 AlbCre mice, key pro-inflammatory cytokines and hepatic immune cell populations were characterized in-depth (Figure 1D-H). In fact, non-hepatocyte cells of LRH-1-deficient livers expressed higher levels of tumor necrosis factor (TNF), interleukin (IL)-1β and IL-6, suggesting an ongoing inflammatory response (Figure 1D). High-dimensional flow cytometry analysis of these cells revealed major differences in immune cell subsets between LRH-1-proficient and LRH-1-deficient livers (Figure 1E-H, S1B-E). The decline of Kupffer cells (KCs), the resident macrophage population in the liver, and the concomitant increase of monocytes and monocyte-derived cells, including dendritic cells (DCs), are hallmarks of ongoing hepatic inflammation (Campana et al., 2021), which was also seen in untreated LRH-1-deficient livers of both sexes (Figure 1F). In addition to the more abundant macrophage and DC populations, also more neutrophils and several lymphoid subsets, such as T cells, natural killer T cells (NKTs) and innate lymphoid cells (ILCs), were significantly increased in LRH-1-deficient livers, in particular in male mice (Figure 1H). The major difference was observed in the lipid-sensing γδ-T cell population, which was strongly increased in LRH-1 AlbCre livers, but barely detectable in healthy control mice (Figure 1H). Taken together, untreated LRH-1 AlbCre mice have a disturbed hepatic lipid metabolism, presumably due to the lack of LRH-1 as a key metabolic regulator in hepatocytes. Our results indicate that this lipid disbalance is involved in recruiting and activating immune cells (Hubler and Kennedy, 2016), ultimately leading to a decline of regulatory KCs, restricted immune cell-mediated hepatocyte death and concomitant activation of collagen-producing stellate cells.

**Figure 1:**
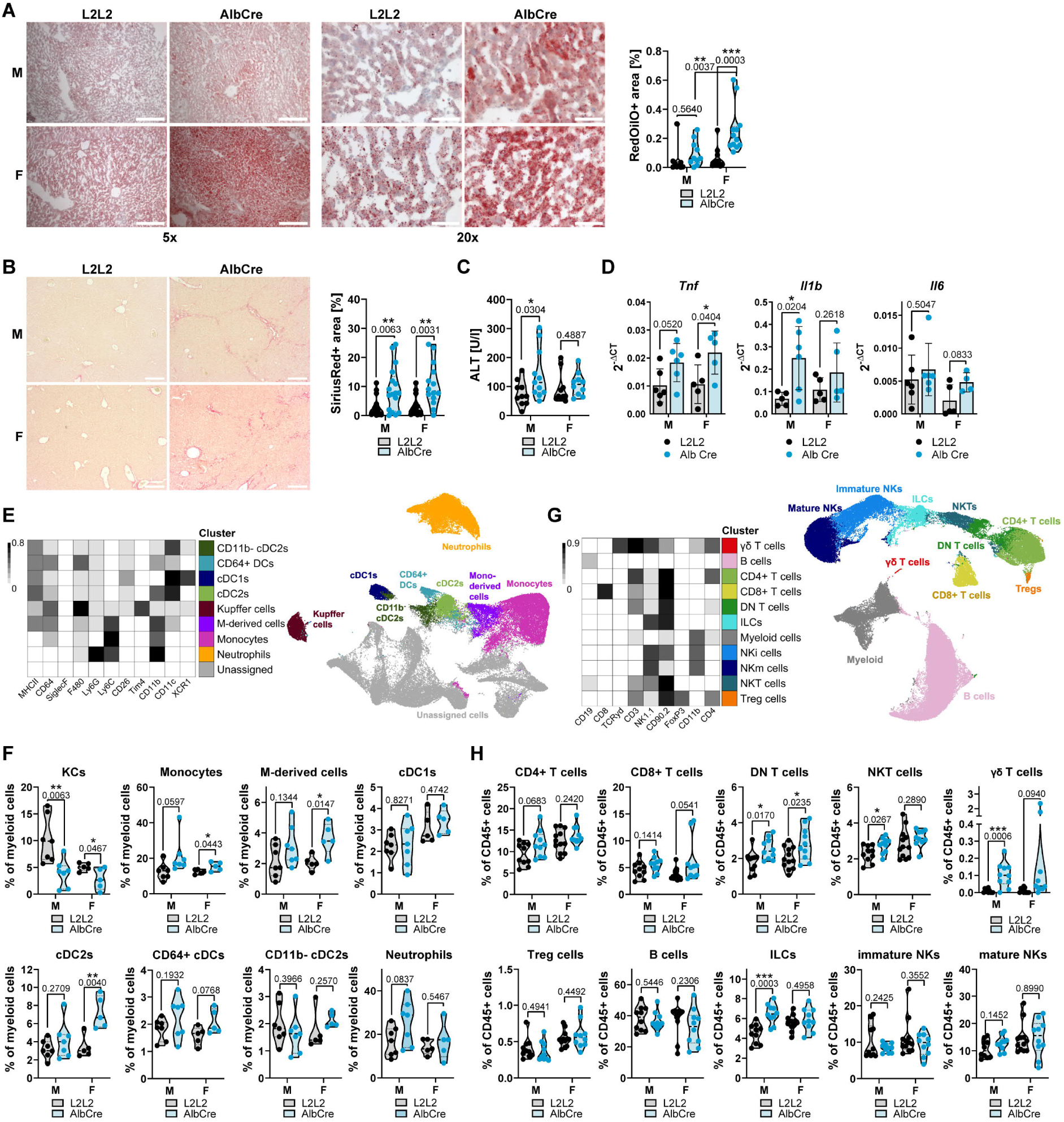
Hepatic LRH-1 deficiency provokes steady-state steatosis and inflammation. Data are from male (M) and female (F) LRH-1 L2L2 (L2L2) and LRH-1 AlbCre (AlbCre) mice. (A) Representative pictures and quantification of OilRedO-stained fat in liver sections. Scale bar 200 µm (left), 50 µm (right). (B) Representative pictures and quantification of SiriusRed-stained collagen in liver sections. Scale bar 250 µm. (C) Serum alanine aminotransferase (ALT) levels of untreated mice. (D) Transcript levels of non-parenchymal liver cells isolated from untreated mice. (E-H) Computational flow cytometry analysis of non-parenchymal liver cells, divided into myeloid (E-F) and lymphoid immune cells (G-H). Populations are identified and visualized using FlowSOM and UMAP algorithm, and annotated based on marker expression shown in heatmaps. KCs: Kupffer cells, M-derived cells: monocyte-derived cells, cDCs: conventional dendritic cells, NKT: natural killer T cells, ILCs: innate lymphoid cells, NKs: natural killer cells. Bar graphs show mean +/- SD and violin plots show median with quartiles. Dots represent values from individual mice. Statistical significance tested by Two-way ANOVA with Sidak’s multiple comparison test.

### Sex-dependent regulation of TNF-induced hepatitis

Next, we investigated next how the described inflammatory phenotype of LRH-1 AlbCre mice influences their sensitivity to different experimental models of hepatitis. Paracetamol (Acetaminophen, APAP) overdose-induced liver failure is a classical model to investigate acute drug-induced liver damage and associated inflammation (Mossanen and Tacke, 2015). In line with recent results (Sun et al., 2023), we found that male LRH-1-deficient mice were more sensitive to APAP-induced liver damage (Figure 2A-B). Notably, APAP toxicity relies to a great extent on the exceptional damage of mitochondria with associated depletion of cellular energy (Ramachandran and Jaeschke, 2019a). We found that both glycolytic and mitochondrial ATP production were strongly attenuated in LRH-1-deficient hepatocytes (Figure 2SA-B). This restricted energy phenotype worsened APAP-provoked ATP depletion and increased the sensitivity towards APAP-induced liver injury (Figure S2B). Interestingly, in contrast to male mice, female LRH-1 AlbCre mice were not more susceptible to APAP toxicity than their female control littermates (Figure 2A), highlighting critical sex-specific differences in APAP-induced liver damage and inflammation.

**Figure 2:**
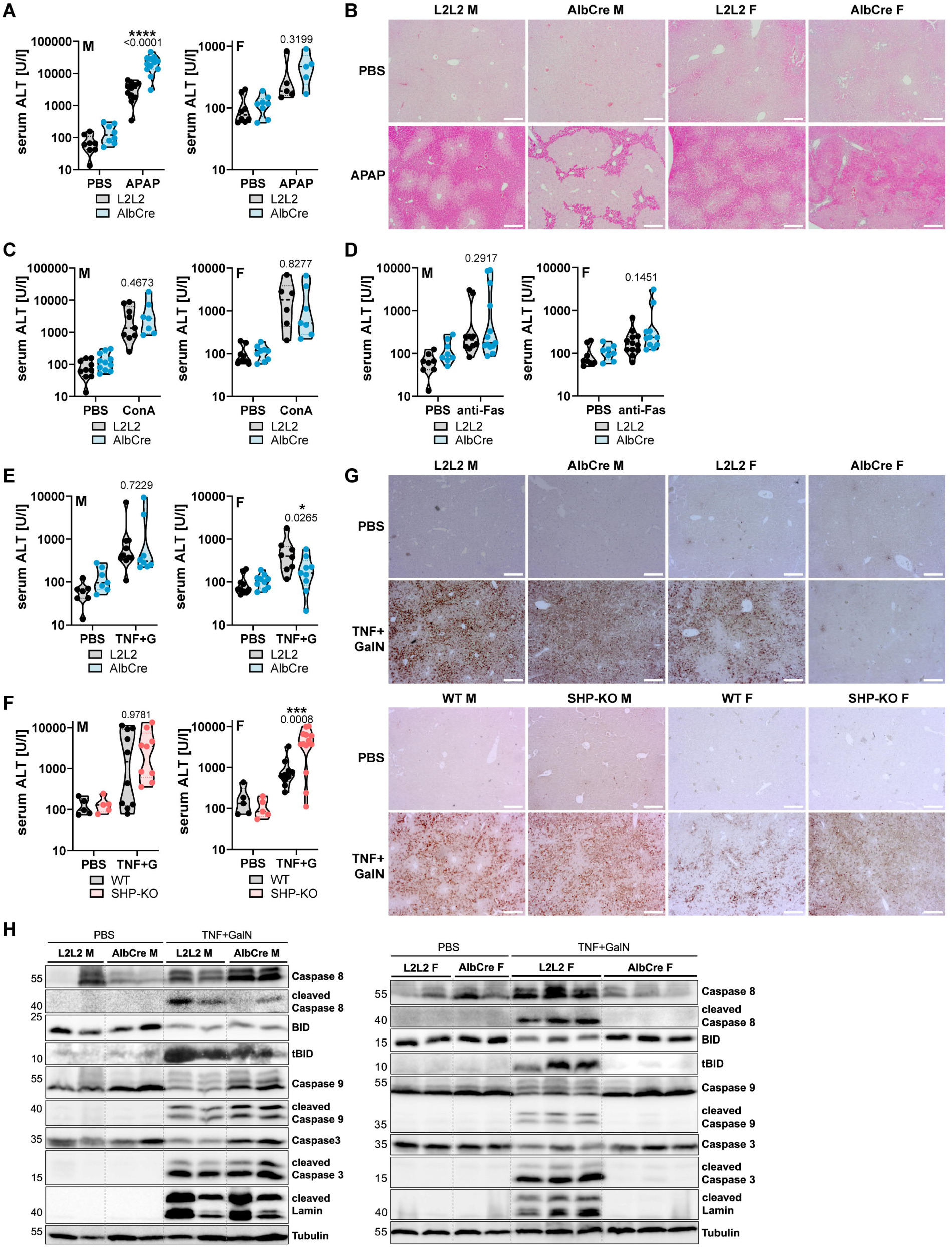
Hepatocyte LRH-1 sex-dependently regulates TNF-induced hepatitis. Data are from male (M) and female (F) LRH-1 L2L2 (L2L2) and LRH-1 AlbCre (AlbCre) mice. (A) Serum alanine aminotransferase (ALT) of mice treated with 500 mg/kg APAP for 6h. (B) Representative pictures of H&E-stained in liver sections of mice described in (A). Scale bar 250 µm. (C-F) Serum ALT of mice treated with 15 mg/kg Concanavalin A (ConA) for 8h (C), 150 µg/kg anti-Fas clone Jo2 for 8h (D), or 25 µg/kg murine TNF and 1000 mg/kg N-Galactosamine (G, GalN) for 6h (E-F). (G) Cleaved Caspase 3 Immunohistochemistry of liver sections of mice treated as described in (E-F). Scale bar 250 µm. (H) Western Blot of liver lysates of mice treated with 25 µg/kg murine TNF and 1000 mg/kg N-Galactosamine (G, GalN) for 6h. Detected proteins are shown on the right, molecular weight is shown in kDa on the left. Violin plots show median with quartiles. Dots represent individual mice. Statistical significance tested by Two-way ANOVA with Sidak’s multiple comparison test.

As a model for immune cell-mediated hepatitis, we studied next Concanavalin A (ConA)-induced acute liver damage (Heymann et al., 2015). The lectin ConA activates T cells, which in turn promote extrinsic apoptosis in hepatocytes in a TNF- and Fas Ligand (FasL/CD95L)-dependent manner (Ksontini et al., 1998). Activation of death receptors, i.e. TNF receptor (TNFR) 1 and Fas, is a general hallmark of inflammatory liver diseases and thus a good model to study physiologically relevant disease conditions. Interestingly, there was no difference between LRH-1-deficient and control animals in the sensitivity to ConA-induced hepatitis (Figure 2C). Also, Fas-induced liver injury was comparable between control and LRH-1-deficient mice (Figure 2D). Strikingly, in contrast to APAP-induced liver injury and opposite to our expectations, we found that female LRH-1-deficient mice were significantly protected from TNF-induced liver injury (Figure 2E). SHP (small heterodimer partner, NR0B2) is a transcriptional target of LRH-1 and represses its activity in a negative feedback loop (Ortlund et al., 2005). Confirming this paradoxical finding, female SHP-deficient mice showed the opposed phenotype to LRH-1-deficient mice, as they were more sensitive to TNF-induced liver damage (Figure 2F-G). Upon ligand-induced activation of the TNFR1, several signaling molecules are recruited, promoting the activation of mitogen-activated protein kinases (MAPK) and nuclear factor kappa-light-chain-enhancer of activated B cells (NF-ⲕB), ultimately leading to pro-inflammatory and pro-survival responses (Holbrook et al., 2019). However, blocking these survival signals, e.g. by the hepatocyte-specific transcription inhibitors D-galactosamine (GalN) or Actinomycin D (ActD), or by interfering with ubiquitination processes at the receptor complex, results in activation of downstream caspases and apoptotic cell death induction (Cockram et al., 2021; Holbrook et al., 2019; Leist et al., 1994). Strikingly, LRH-1-deficient female mice did not show any of the typical processing of Caspase 8, BID, Caspase 3 or its targets in response to TNF and GalN treatment (Figure 2H). Notably, LRH-1 deficiency protected also from bacterial lipopolysaccharide (LPS) and GalN-induced hepatitis (Figure S2C-E). LPS activates hepatic macrophages that produce TNF, which in turn kills hepatocytes in the presence of GalN. Although serum TNF levels were reduced in LPS-or TNF-treated LRH-1 AlbCre mice compared to treated control mice (Figure S2F), hepatic macrophage activity was not compromised as isolated hepatic macrophages of both genotypes were able to produce TNF in response to LPS stimulation (Figure S2G). Collectively, our results indicate that the critical factor mediating the protection from TNF-induced liver damage in LRH-1 AlbCre mice must lay within the TNF-induced cell death pathway in hepatocytes.

### LRH-1-deficient hepatocytes are protected from TNF-induced cell death

To investigate the underlying mechanism of the protection observed *in vivo*, we treated hepatocytes isolated from LRH-1 AlbCre and control mice with TNF and Actinomycin D (ActD). Surprisingly, LRH-1-deficient hepatocytes from both sexes were protected from TNF-induced cell death (Figure 3A-C), while no protection was observed when treating hepatocytes with APAP or cisplatin (Figure 3D). Hepatocytes are defined as Type II cells, in which death receptor-induced apoptosis requires the amplification via the mitochondrial apoptosis pathway with a critical contribution of the BCL-2 homologs BIM and BID (Corazza et al., 2006; Jost et al., 2009; Kaufmann et al., 2009). Activation of these two pro-apoptotic BCL-2 homologs results in mitochondrial outer membrane permeabilization (MOMP), which is required for downstream caspase activation. MOMP was strongly attenuated in TNF/ActD-treated LRH-1-deficient hepatocytes compared to control hepatocytes, as seen by the reduced release of cytochrome c and SMAC from mitochondria into the cytosol (Figure 3E). This suggests that LRH-1 deficiency affects TNF-induced cell death upstream of MOMP, for example at the TNFR1 complex or at the level of the BCL-2 interactome.

**Figure 3:**
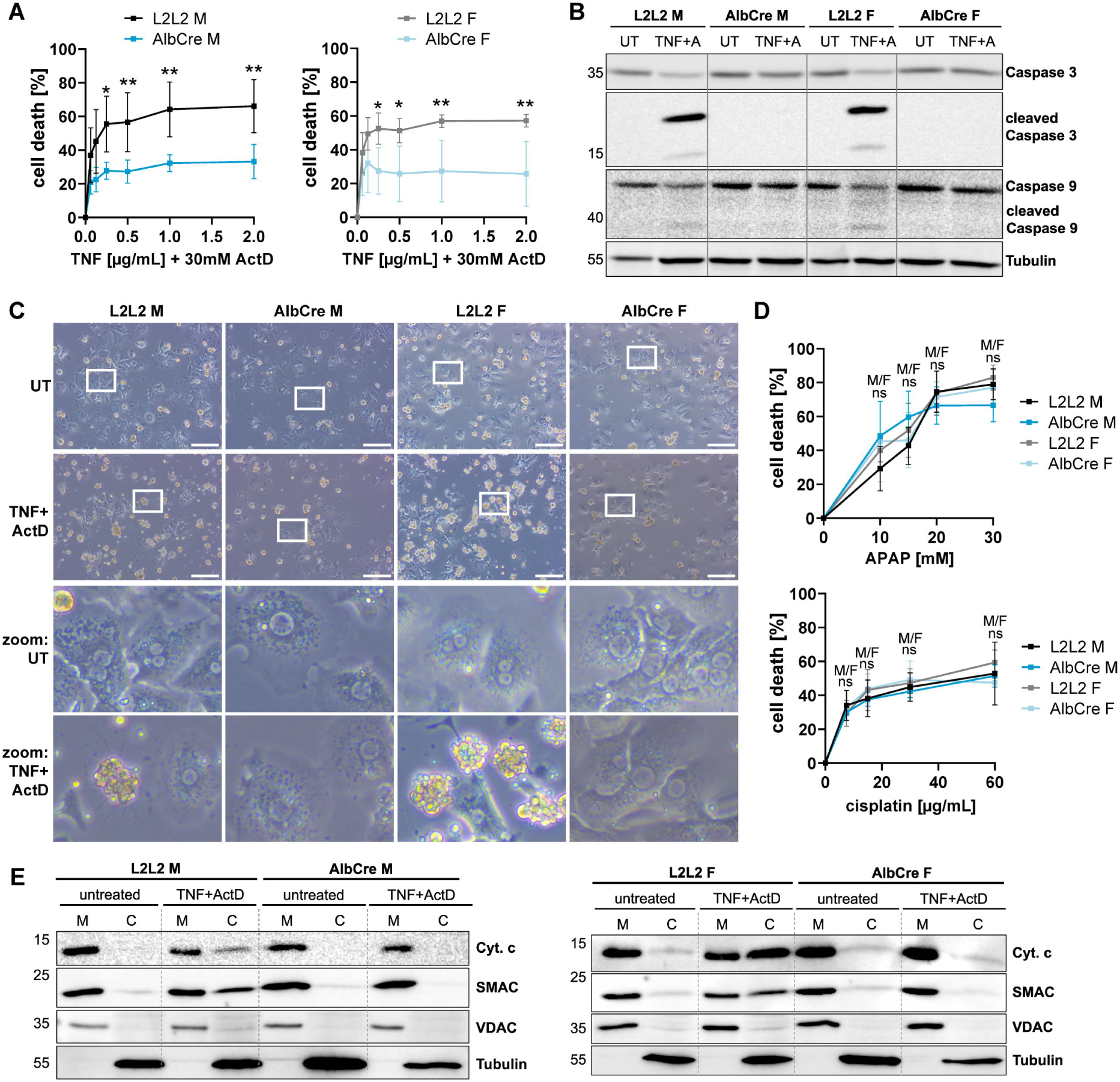
LRH-1-deficient hepatocytes are protected against TNF-induced cell death. Data are from primary hepatocytes of male (M) and female (F) LRH-1 L2L2 (L2L2) and LRH-1 AlbCre (AlbCre) mice. (A) MTT cell death assay of hepatocytes treated with TNF and Actinomycin D (ActD, A) for 16h. n=4-6. (B) Western blot of hepatocytes treated with 2 ng/ml murine TNF and 30 nM ActD for 12h. Detected proteins are shown on the right, molecular weight is shown in kDa on the left. (C) Representative pictures of hepatocytes treated with 2 ng/ml murine TNF and 30 nM ActD for 8h. Scale bar 150 µm. (D) MTT cell death assay of hepatocytes treated with APAP or cisplatin for 16h. n=4-8 (APAP), n= 3-7 (cisplatin). (E) Western blot of mitochondrial (M) and cytosolic (C) fractions of hepatocytes treated with 2 ng/ml murine TNF and 30 nM ActD for 6h. Detected proteins are shown on the right, molecular weight is shown in kDa on the left. Line graphs show mean +/- SD with n as independent experiments. Statistical significance tested by Two-way ANOVA with Sidak’s multiple comparison test.

Interestingly, the *ex vivo* results do not fully match those of *in vivo* experiments, as sex-dependent differences in TNF sensitivity were only observed *in vivo*. This may be explained by the multifaceted roles of LRH-1 and by the more complex hepatic environment *in vivo* compared to isolated hepatocytes. In line with differences in hepatic immune cell populations between male and female LRH-1-deficient mice (Figure 1), expression of pro- and anti-inflammatory cytokines in the liver varied between both sexes (Figure S3A). Although most pro-inflammatory cytokines were increased in LRH-1 AlbCre livers of both sexes, TNF-induced IL-6 was elevated in male LRH-1-deficient livers, but significantly reduced in female livers (Figure S3A). Also, LRH-1 transcriptionally regulates the expression of several steroidogenic enzymes (Mueller et al., 2006; Sirianni et al., 2002). In the liver, the steroidogenic enzyme 11β-hydroxysteroid dehydrogenase (*Hsd11b1*) is critical for the production of anti-inflammatory glucocorticoids (Semjonous et al., 2011) and glucocorticoids in the liver were actually shown to cause sex-dependent differences in gene expression determining the sensitivity to inflammation (Duma et al., 2010). Interestingly, female LRH-1 AlbCre mice showed lower hepatic expression of this enzyme, however, actual glucocorticoid levels were unaffected in liver and serum (Figure S3B-D). The sex differences between LRH-1 AlbCre mice in hepatitis might also be attributed to an altered abundancy of sex steroid hormones, since androgen and estrogen receptors were shown to critically modulate inflammatory processes (Mohamad et al., 2019; Straub, 2007) and as LRH-1 is heavily intertwined in their abundancy and activity (Annicotte et al., 2005; Lai et al., 2013; Thiruchelvam et al., 2011; Zhou et al., 2005). Androgen- and estrogen-metabolizing enzymes were differentially expressed among both genotypes and sexes, overall indicating less sex hormones in LRH-1-deficient livers (Figure S3E-F). Thus, the deletion of LRH-1 in the liver has likely wide-spread and complex effects, which may explain the difference between *in vivo* and *in vitro,* and between male and female mice.

### LRH-1 inhibits NF-kB activity by direct interaction

As LRH-1 deficiency-associated protection was restricted to TNF-induced cell death, we focused next on TNFR1 signaling. Ubiquitination of TNFR signaling complex proteins defines whether downstream signals proceed into Caspase 8-dependent apoptosis or NF-ⲕB-mediated survival (Cockram et al., 2021; Figure S4A). Ubiquitination of TNFR1 complex proteins is mediated in part by cellular inhibitor of apoptosis proteins (cIAP) 1 and cIAP2, and removed by the deubiquitinases Otulin, A20 and CYLD (Cockram et al., 2021). While cIAP1 and cIAP2 expression was not altered, Otulin and A20 were reduced in LRH-1-deficient livers, suggesting enhanced pro-survival signaling (Figure S4B-C). However, the pattern of IⲕB degradation and MAPK phosphorylation did not differ between LRH-1-deficient and control hepatocytes (Figure S4D), indicating that TNFR1 signaling is rather not critically altered.

Interestingly, we found that in LRH-1 AlbCre hepatocytes the NF-ⲕB subunit RelA, which in resting cells is usually cytosolic but translocates to the nucleus upon cellular activation, accumulated in the nucleus, even in the absence of TNF stimulation (Figure 4A). In line with increased nuclear accumulation of RelA, NF-ⲕB activity was significantly enhanced in LRH-1 deficient hepatocytes (Figure 4B). By employing an inducible LRH-1 overexpression in a hepatocyte cell line, we identified that increased LRH-1 expression and activity negatively correlated with NF-ⲕB activity (Figure 4C) *Vice versa*, TNF-induced increase in NF-ⲕB activity lowered LRH-1 activity (S4E-G). LRH-1 overexpressing cells appeared also to be slightly more sensitive to TNF-induced cell death (Figure S4F). Since diminished NF-ⲕB activity in LRH-1-overexpressing cells may be caused by their previously described mutual interaction (Huang et al., 2014), we confirmed that LRH-1 and RelA can be co-precipitated (Figure 4D). To investigate this interaction in more depth, we analyzed binding of LRH-1 and RelA by microscale thermophoresis, employing purified full length LRH-1 or the ligand-binding domain (LBD) of LRH-1, and three different peptides of RelA (Figure 4E). All combinations showed binding and reasonable dissociation constants (Kd). Thus, our results identified that the LBD of LRH-1 directly interacts with the Rel homology domain (RHD) of RelA (Figure 4F, S4H), which solidifies the regulatory role of LRH-1 in TNF signaling by modulating NF-ⲕB activity.

**Figure 4:**
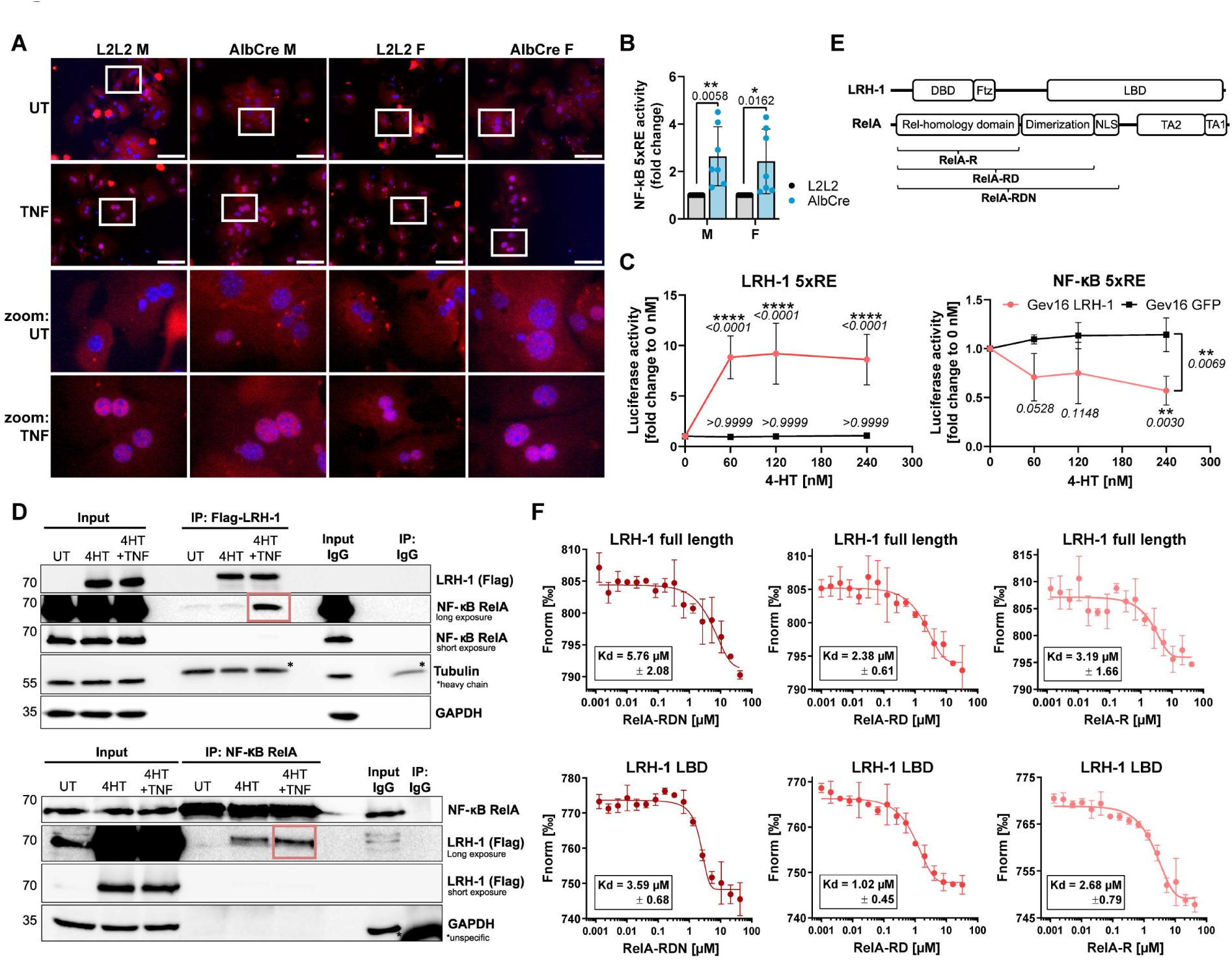
LRH-1 inhibits NF-ⲕB activity by direct interaction. (A) Representative immunocytochemistry pictures of hepatocytes of male (M) and female (F) LRH-1 L2L2 (L2L2) and LRH-1 AlbCre (AlbCre) mice. Hepatocytes were treated with 10 ng/ml murine TNF for 1h, fixed and stained with anti-RelA antibody (red) and DAPI (blue). Scale bar 100 µm. (B) NF-ⲕB activity measured by luciferase assay in hepatocytes of mice described in (A). Untreated hepatocytes were transfected with plasmids containing luciferase gene behind 5x response element (5xRE) of NF-ⲕB and analyzed after 24h. (C) LRH-1 and NF-ⲕB activity measured by luciferase assay in IHH Gev16 LRH-1 or GFP cells that have 4-hydroxytamoxifen (4-HT)-inducible Flag-tagged human LRH-1 overexpression or GFP overexpression. IHH cells were treated with 4-HT for 24h, then transfected with plasmids containing luciferase gene behind 5xRE of LRH-1 (left) or NF-ⲕB (right) and analyzed after 24h (n=6). (D) Western Blot of Immunoprecipitation of Flag (up) and RelA (low) in IHH Gev16 LRH-1 cells (described in C). IHH cells were treated with 60 nM 4-HT for 24h, treated with 10 ng/ml human TNF for 1h and lysed. Boxes indicate binding of LRH-1 and RelA. Detected proteins are shown on the right, molecular weight is shown in kDa on the left. (E) Peptides used for experiments in (F): Full length LRH-1 peptide or only Ligand binding domain (LBD) peptide of LRH-1; three different RelA peptides: RelA-R, RelA-RD and RelA-RDN. DBD: DNA-binding domain; NLS: nuclear localization domain; TA: transactivation domain. (F) Microscale Thermophoresis binding curves and dissociation constants (Kd) of peptides shown in (E) (n=3 each). Bar plots and line graphs show mean +/- SD with n as independent mouse (B) or experiments (C, F). Statistical significance tested by Two-way ANOVA with Sidak’s multiple comparison test.

### Hepatocyte-specific LRH-1 deficiency upregulates anti-apoptotic BCL-2 proteins

Increased NF-ⲕB activity in LRH-1-deficient hepatocytes could explain the observed increased inflammatory signaling (Figure 1) and the shift towards pro-survival responses upon TNF treatment (Figure 2 and 3). NF-ⲕB transcriptionally regulates several anti-apoptotic proteins, including cIAP1/2 and BCL-2-like proteins (Chen et al., 2000; Karin and Lin, 2002). While cIAP1 and 2 expression was not altered (Figure S4B), we found that BCL-2 and MCL-1 transcript levels were upregulated in LRH-1-deficient livers, and *vice versa* downregulated in LRH-1-overexpressing cells (Figure 5A-B). On a protein level, LRH-1 deficiency clearly resulted in higher BCL-2 and MCL-1 expression in untreated livers and hepatocytes (Figure 5C-D, S5A). More abundant anti-apoptotic BCL-2 homologs could neutralize more pro-apoptotic BCL-2 proteins, and hence would prevent MOMP and cell death in LRH-1 AlbCre hepatocytes upon TNF treatment (Figure 3E).

**Figure 5:**
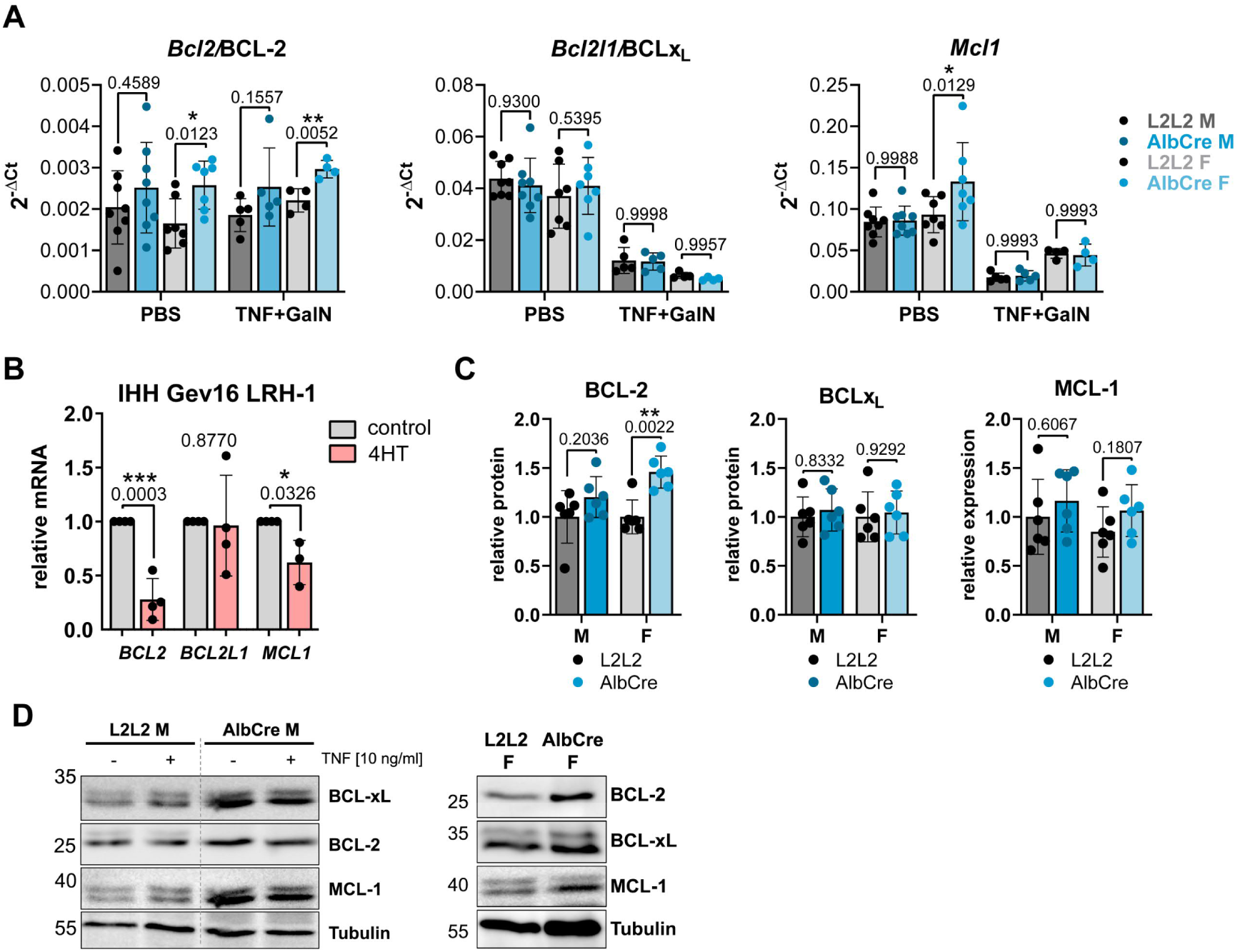
Anti-apoptotic BCL-2-like proteins are upregulated in LRH-1 AlbCre hepatocytes. (A) Transcript levels of liver samples from male (M) and female (F) LRH-1 L2L2 (L2L2) and LRH-1 AlbCre (AlbCre) mice treated with 25 µg/kg murine TNF and 1000 mg/kg N-Galactosamine (G, GalN) for 6h. (B) Transcript levels of IHH Gev16 LRH-1 cells with treated with 60 nM 4-hydroxytamoxien (4-HT) for 24h. (C) Quantification of protein expression based on western blots shown in Figure S5. Expression was calculated by comparison to loading control and normalized to mean of L2L2 mice of each sex. (D) Western Blot of hepatocytes untreated or treated with murine TNF for 2h. Detected proteins are shown on the right, molecular weight is shown in kDa on the left. Bar graphs show mean +/- SD with each dot as independent mouse (A, C) or experiment (B). Statistical significance tested by Two-way ANOVA with Sidak’s multiple comparison test (A, C) or by unpaired student’s t test (B).

### Hepatic regeneration causes degradation of pro-apoptotic BIM in LRH-1 AlbCre mice

In Type II cells, TNF-induced apoptosis proceeds via the pro-apoptotic proteins BID and BIM and amplification via the mitochondrial apoptosis pathway. While BID is cleaved and activated by Caspase 8, BIM becomes phosphorylated and activated by c-Jun kinase (JNK). Consequently, BID- and BIM-deficient mice show reduced sensitivity to TNF-induced hepatitis (Jost et al., 2009; Kaufmann et al., 2009). Accordingly, TNF treatment resulted in strong phosphorylation of JNK and BIM in control mice and less pronounced also in LRH-1 AlbCre mice (Figure 6A, S6A). While BIM transcript levels were unaffected (Figure 6B), BIM proteins were remarkably reduced in untreated, LRH-1-deficient livers and hepatocytes (Figure 6C-E, S6B-C). Reduced BIM levels are decisive for the outcome of TNF-induced hepatocyte death as BIM-deficient mice are protected from LPS/GalN-induced (TNF-mediated) liver damage (Figure 6F). Since reduced mRNA expression is likely not the underlying reason for low BIM protein levels, we investigated whether increased proteasomal degradation may be responsible for it. In support of this notion, our results indeed demonstrated that BIM protein levels in LRH-1-deficient hepatocytes could be completely restored when treated with the proteasome inhibitor MG132 (Figure 6G). Proteasomal degradation of BIM is initiated by ERK1/2-mediated phosphorylation at distinct serine sites (Luciano et al., 2003) and subsequent ubiquitylation-mediated proteasomal degradation. ERK1/2 is typically activated in response to growth factor signaling, and we found several hepatic growth factors being upregulated in LRH-1-deficient livers (Figure 6H). Growth factors indicate ongoing regeneration in response to damage, as illustrated by the inflammatory and fibrotic phenotype of LRH-1 AlbCre livers (Figure 1). Strikingly, pharmacological inhibition of growth factor signaling by Tyrphostin, an inhibitor of receptor tyrosine kinases, not only restored BIM levels in LRH-1-deficient hepatocytes, but also re-sensitized them to TNF-induced cell death (Figure 6I-J). Overall, these results demonstrate that hepatic LRH-1 critically orchestrates hepatic cell death and hepatitis as the liver-specific deletion of LRH-1 provokes inflammation by lipid deposition and increased NF-ⲕB activity, and at the same time prevents cell death by NF-ⲕB-mediated pro-survival signaling and growth factor-driven degradation of BIM.

**Figure 6:**
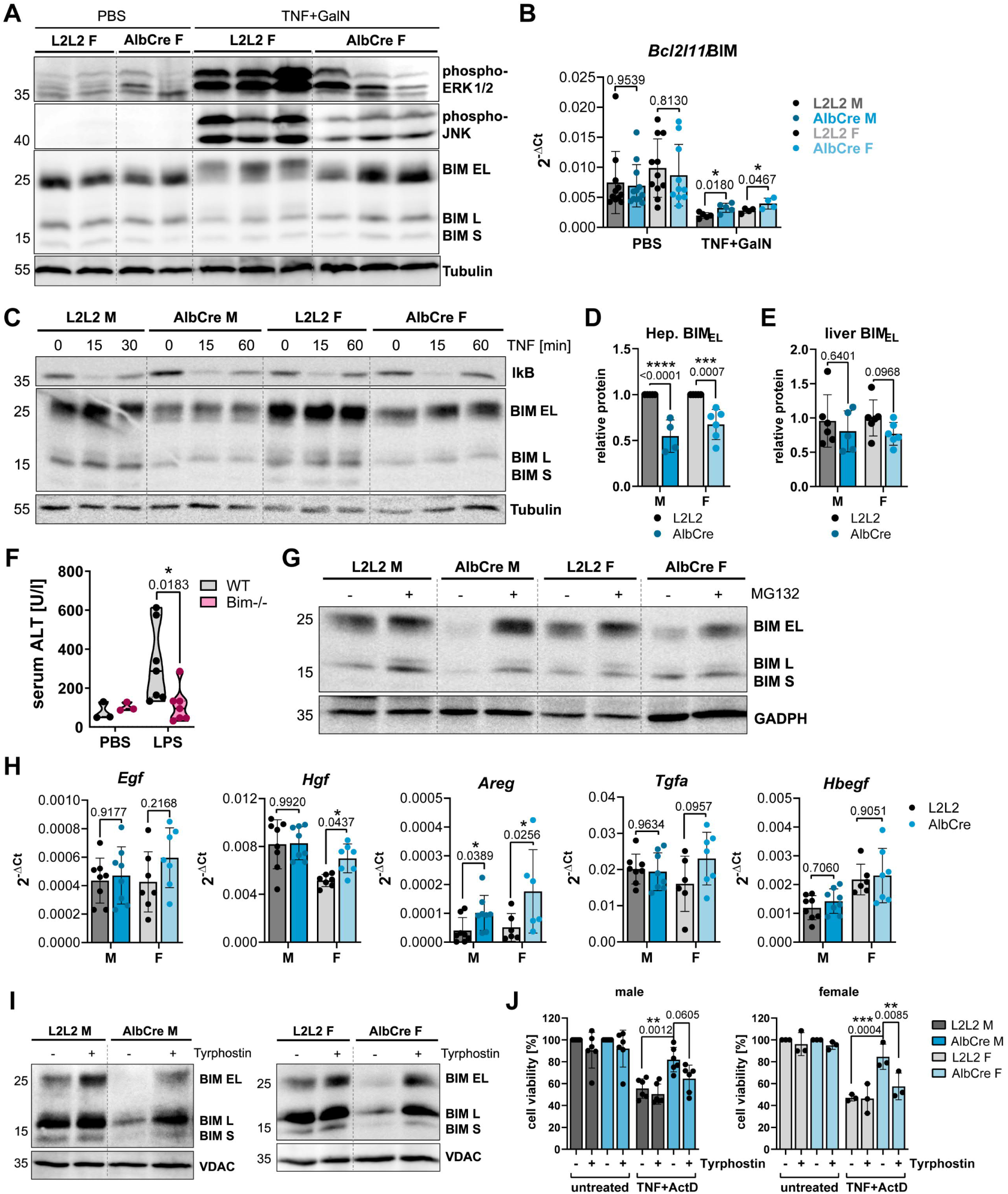
Hepatic regeneration causes degradation of pro-apoptotic BIM in LRH-1 AlbCre mice. Data are from primary hepatocytes of male (M) and female (F) LRH-1 L2L2 (L2L2) and LRH-1 AlbCre (AlbCre) mice. (A) Western Blot of liver lysates of mice treated with 25 µg/kg murine TNF and 1000 mg/kg N-Galactosamine (G, GalN) for 6h. (B) Transcript levels of liver from mice described in (A). (C) Western Blot of hepatocytes treated for indicated time with 10 ng/ml TNF. (D-E) Quantification of protein expression in hepatocytes (D) and liver lysates (E) based on western blots shown in Figure S6. Expression was calculated by comparison to loading control and then normalized to L2L2 hepatocytes in each blot (D) or to mean of all L2L2 livers of each sex (E). Only untreated samples were considered for quantification. (F) Serum alanine aminotransferase (ALT) levels of wild type (WT) and BIM-/-mice treated with 5 µg/kg LPS and 1000 mg/kg N-Galactosamine for 6h. (G) Western blot of hepatocytes treated with 5 µM MG132 (proteosome inhibitor) for 6h. (H) Transcript levels of liver from untreated mice. (I) Western Blot of hepatocytes treated with 50 µM Tyrphostin for 6h. (J) MTT cell death assay of hepatocytes treated with 50 µM Tyrphostin for 6h and then with 2 ng/ml murine TNF and 30 nM Actinomycin D (ActD) for 16h. Western Blots show detected proteins on the right and molecular weight as kDa on the left. Bar graphs show mean +/- SD and violin plots show median with quartiles. Dots are independent mice. Statistical significance tested by Two-way ANOVA with Sidak’s multiple comparison test.

## Discussion

Here, we report about a paradox role of LRH-1 in regulating liver inflammation and cell death. In line with the previously described role of LRH-1 in regulating lipid metabolism, we found that liver-specific deletion of LRH-1 causes accumulation of lipid droplets, inflammation and associated immune cell infiltration, resulting in mild fibrosis. The increased collagen deposition represents the cause of increased tissue damage and regeneration, as illustrated by increased ALT levels and elevated expression of EGF receptor ligands. Surprisingly, despite signs of liver damage in LRH-1-deleted livers, and a previously reported anti-apoptotic role of LRH-1, we observed that LRH-1 deficiency strongly protected from TNF-induced liver damage and hepatocyte death. Our data suggest that the absence of LRH-1 results in increased NF-ⲕB activation, expression of pro-inflammatory cytokines and induction of pro-survival factors, such as anti-apoptotic members of the BCL-2 family. The elevated NF-ⲕB activation is the result of missing physical interaction with LRH-1 and associated transcriptional inhibition. Physical interaction between RelA and LRH-1 had been reported previously in the context of intestinal inflammation (Huang et al., 2014) and could be confirmed in this study. The presented biochemical studies revealed that the ligand-binding domain (LBD) directly interacts with the Rel homology domain (RHD) of RelA, resulting in mutual inhibition of both transcription factors. Thus, absence of hepatic LRH-1 increases basal NF-ⲕB activity and associated hepatic inflammation, but likewise enhanced resilience towards cell death by elevated expression of anti-apoptotic proteins. The low level of inflammation is likely the driver of regenerative processes including the here reported elevated expression of growth factors. We demonstrate that the EGF receptor signaling-mediated degradation of the pro-apoptotic BCL-2 homolog BIM is critical for the survival of LRH-1-deficient hepatocytes, as pharmacological inhibition of EGF receptor activation re-sensitized LRH-1-deficient hepatocytes to TNF-induced cell death.

Previous studies proposed an anti-apoptotic and anti-inflammatory role for LRH-1 in various tissues and disease contexts. Loss of local LRH-1 was shown to promote intestinal and pancreatic inflammation, and cell death in enterocytes and β-islet cells, respectively (Bayrer et al., 2018; Cobo-Vuilleumier et al., 2018b; Coste et al., 2007; Martin Vázquez et al., 2022; Mueller et al., 2006). In line, we report here that hepatic LRH-1 deletion sensitized to APAP-induced liver injury. Accordingly, our finding that LRH-1 deficiency protects from TNF-induced hepatitis was rather surprising. These opposing effects of LRH-1 deletion on TNF- and APAP-induced liver damage can be explained by their different mechanisms of toxicity. APAP overdose drastically depletes cellular glutathione and ATP levels, provoking hepatocyte necrosis and rapidly activating the hepatic immune system. In contrast, TNF-induced apoptosis is a rather non-immunogenic, ‘silent’ form of cell death. Additionally, it can be intracellularly modulated by the apoptotic signaling machinery. Hence, the pro-inflammatory milieu in LRH-1-deficient livers presumably amplifies APAP-induced liver damage but has only little effect on TNF-induced apoptosis, and *vice versa*, the elevated NF-ⲕB activity and associated expression of anti-apoptotic BCL-2 homologs counteract TNF-induced apoptosis, but likely does not affect the APAP-induced cellular collapse. Thus, the role of LRH-1 in the modulation of inflammation and cell death is highly disease-dependent. Yet, also within the same pathological condition, LRH-1 can play opposing roles depending on the cell type examined. We report here that hepatocyte LRH-1 deficiency protects from TNF- and LPS-induced hepatitis via cell-intrinsic pro-inflammatory and pro-survival NF-ⲕB signaling. However, in a previous study, our group demonstrated that pharmacological inhibition of LRH-1 protects from LPS-induced hepatitis by limiting pro-inflammatory functions of macrophages and associated cytokine release (Schwaderer et al., 2020). These opposing cell type-dependent actions of LRH-1 on inflammation can be explained by different LRH-1 expression levels in tissue cells, with high LRH-1 expression, and cells of the hematopoietic system, with low expression. It was shown that transcription factor abundancy dose-dependently determines gene expression (Brewster et al., 2014) modulating downstream effects. Additionally, different levels of LRH-1 will also affect its interaction with other transcription factors, which will define cell identity and response to stress (Feng et al., 2023; Joo et al., 2019). Consequently, the disease mechanism and the involved cell types need to be carefully considered when examining and manipulating LRH-1-regulated processes.

Our study furthermore suggests that LRH-1 acts in a sex-dependent manner as the sensitization to APAP intoxication and the protection from TNF-induced hepatitis were only observed in male, respectively female mice. Among other factors, male and female organisms differ in their sex hormone abundancy, which is, interestingly, regulated by LRH-1 through the expression of cytochrome P450 enzymes involved in steroid hormone synthesis or degradation (Dubé et al., 2009; Sirianni et al., 2002; Xiao et al., 2018; Zhou et al., 2021). Our expression analysis suggests that the levels of both estrogen- and androgen-metabolizing enzymes are reduced in LRH-1-deficient livers. This fits with previous studies establishing a positive correlation between LRH-1 and androgen and estrogen abundancy (Lazarus et al., 2012; Meinsohn et al., 2019; Zhou et al., 2005). In addition to hormone synthesis, LRH-1 can regulate estrogen receptor (ER) activity via two additional mechanisms. Firstly, LRH-1 is a co-activator of the ER as both receptors bind together to specific ER response elements enhancing the respective target gene expression (Chand et al., 2012) and secondly, both transcription factors were shown to mutually regulate their expression (Annicotte et al., 2005; Thiruchelvam et al., 2011). As estrogen and androgen derivatives were shown to critically affect sensitivity to inflammatory diseases, including hepatitis (Straub, 2007), hepatic LRH-1 deficiency has the potential to modulate these effects in a sex-dependent manner. Interestingly, ER activity is believed to regulate anti-inflammatory processes particularly by perturbing NF-ⲕB activity (Gionet et al., 2009). Thus, in addition to the lack of inhibitory interaction with LRH-1, elevated NF-ⲕB activity in LRH-1-deficient hepatocytes might be also attributed to less active ER. It is conceivable that female LRH-1 AlbCre mice are more affected by the proposed reduction in ER activity than their male littermates and consequently exhibit a stronger inflammatory phenotype leading to protection from hepatitis. In line, female mice showed a more pronounced increase in anti-apoptotic BCL-2 proteins, suggesting that female mice could have elevated apoptotic resistance in terms of mitochondrial priming. Mitochondrial priming describes the susceptibility of cells to undergo apoptosis based on their dependency and availability of specific BCL-2 homologs (Potter and Letai, 2016). Female LRH-1-deficient mice are presumably less susceptible due to the described dual potentiation of NF-ⲕB activity and associated expression of BCL-2 homologs. Nonetheless, it is important to note that *in vitro* LRH-1-deficient hepatocytes from both male and female mice were found to be resistant to TNF induced apoptosis, supporting the view that the observed resistance is a general phenomenon.

Death receptor-induced hepatocyte apoptosis is dependent on the BH3-only proteins BIM and BID (Corazza et al., 2006; Kaufmann et al., 2009, 2012). LRH-1 deficiency resulted in a substantial reduction in BIM protein levels, which critically contributed to the protection from TNF-induced hepatitis. As the degree of Fas-induced liver damage was rather small, it cannot be excluded that LRH-1 may regulate this process as well. We previously described that BIM deficiency protects from APAP-induced liver damage (Badmann et al., 2011), however, here LRH-1 deficiency with the described decline in BIM exacerbated APAP-induced injury. We assume that particularly the compromised energy metabolism and the pro-inflammatory milieu in LRH-1-deficient mice make them more susceptible to the ATP-depleting APAP treatment (Ramachandran and Jaeschke, 2019a) and later more prone to inflammation in response to the highly immunogenic hepatocyte necrosis (Ramachandran and Jaeschke, 2019b; Wang et al., 2022). The reduction of BIM levels in LRH-1-deficient hepatocytes is likely not sufficient to counteract these detrimental effects.

Intriguingly, hepatocyte-restricted deletion of LRH-1 appears to reshape the entire hepatic environment. Joo *et al*. described that LRH-1 maintains hepatocyte identity and sensitivity to hepatotoxins by regulating key transcription factor circuits in the liver (Joo et al., 2019). It becomes increasingly evident that LRH-1 functions to a great extent via the regulation of several other transcription factors, either by direct interaction or by expression of transcription factors or their ligands (Chong et al., 2012; Dubé et al., 2009; Huang et al., 2014; Joo et al., 2019; Lu et al., 2000; Meinsohn et al., 2019; Michalek et al., 2022; Thiruchelvam et al., 2011; Xiao et al., 2018). This overall suggests an intertwined network of transcription factors with LRH-1 as a central hub. Emerging research stresses that particularly in the liver transcription factor networks cooperate to maintain zonation-dependent functions and to prevent loss of identity and dedifferentiation, as it for example happens during fibrotic steatohepatitis, hepatocellular carcinoma, hypoxia or drug-induced liver injury (Ben-Moshe et al., 2022; Kietzmann, 2019; Loft et al., 2021, 2022; Sia et al., 2017; Walesky et al., 2020). The impact of specific nuclear receptors during acute liver injury was just recently highlighted (Klatt et al., 2023). We propose that chronic lack of LRH-1 disturbs these transcription factor networks and will thus, in long term, reshape the liver environment in a multifaceted manner, including non-hepatocyte cells. Our study highlights the complex role of LRH-1 in the liver as we report that hepatocyte LRH-1 deficiency on one side leads to increased hepatic stress and inflammation, but on the other side protects from TNF-induced liver damage.

## Limitations of the study

Our study characterized the effect of hepatic LRH-1 deletion on the steady-state condition and on experimentally induced hepatitis, and thereby identified critical LRH-1-regulated processes. Although we describe distinct intracellular mechanisms, there is likely more to elucidate considering LRH-1 as critical hub within the hepatic transcription factor network. In addition, we describe remarkable differences in the effect of LRH-1 deficiency between male and female mice, but our analyses failed to fully elucidate the underlying cause. LRH-1 appears to be important for several sex-dependent processes in non-genital organs, like the liver. Thus, in-depth characterization of hepatic hormones, metabolites and transcription factor activities in LRH-1-deficient male and female mice may help to understand general mechanisms. Nonetheless, our study emphasizes to increase awareness on sex differences in biomedical sciences and to include both sexes in given studies.

## Abbreviations

ActD: Actinomycin D
APAP: Paracetamol, Acetaminophen
BCL-2: B-cell lymphoma protein 2
BID: BH3 interacting-domain death agonist
BIM: BCL-2 Interacting Mediator of cell death
cIAP: cellular inhibitor of apoptosis protein
ConA: Concanavalin A
DCs: dendritic cells
F: Female
FasL: Fas (CD95) Ligand
GalN: N-Galactosamine
I B: NF-B inhibitor protein
ILCs: innate lymphoid cells
KCs: Kupffer cells
LBD: ligand binding domain
LPS: Lipopolysaccharide
LRH-1: Liver Receptor Homolog 1
M: Male
MAPK: mitogen-activated protein kinase
MOMP: mitochondrial outer membrane permeabilization
NF-B: nuclear factor kappa-light-chain-enhancer of activated B cells
NKTs: natural killer T cells
RHD: Rel homology domain
SHP: small heterodimer partner
SMAC: second mitochondria-derived activator of caspases
TNF: tumor necrosis factor
TNFR: TNF Receptor

## Acknowledgement

We would like to thank the flow cytometry facility (FlowKon) and the animal facility of the University of Konstanz for technical support and animal caretaking. The LRH-1 AlbCre and SHP-deficient mice were kindly provided by K. Schoonjans (EPFL Lausanne).

## Author contributions (CRediT)

Conceptualization: R.L. and T.B.; Investigation: R.L., M.E.D., V.G., K.S., B.F. and J.F.; Formal Analysis: R.L., A.P.P. and T.S.P.; Writing – Original Draft: R.L.; Writing – Review & Editing:

R.L. and T.B.; Resources: O.M. and T.B.; Funding Acquisition: M.E.D. & B.F. and T.B.; Supervision: T.B.

## STAR Methods

### Animal experimentation

All procedures for animal experiments were approved according to European Union guidelines by the authorities of the state of Baden-Württemberg, Germany. Mice were housed in groups of 2-6 animals in IVC cages in 12:12 h light/dark cycle at central animal facility of the University of Konstanz and had permanent access to water and standard chow ad libitum. Mice with floxed LRH-1 gene and heterozygous Cre recombinase gene under Albumin promoter were a generous gift from Kristina Schoonjans (EPFL Lausanne) (Mataki et al., 2007). For experiments we used littermates that have floxed LRH-1 gene and either no Cre expression, termed “LRH-1 L2L2”, or with hepatic Cre expression, termed “LRH-1 AlbCre”. C57BL/6 mice (wild type mice) were originally obtained from Jackson Laboratory and served as control animals for SHP- and BIM-deficient mice. Mice with systemic SHP deletion were also a kind gift from Kristina Schoonjans (Volle et al., 2007). BIM-deficient (“Bim-/-“) mice mice were a generous gift from Andreas Strasser (Walter and Eliza Hall Institute, Melbourne) (Bouillet et al., 1999). All mice are bred in-house and had been backcrossed to the C57BL/6 background for over more than 10 generations.

For all *in vivo* experiments 8-10 weeks old littermate mice were used. For APAP experiments, mice were fasted for 12h before intraperitoneal (i.p.) injection of 500 mg/kg APAP (Sigma) in warm PBS for 6h (Mossanen and Tacke, 2015). For ConA-induced hepatitis, mice were injected in the tail vein (i.v.) with 15 mg/kg ConA from *Canavalia ensiformis* (Sigma) in PBS for 8 h (Heymann et al., 2015). For Fas-mediated hepatitis, mice were i.p. injected with 150 µg/kg hamster anti-mouse CD95/Fas clone Jo2 antibody in PBS (BD Pharmigen) for 8h. For TNF- and LPS-induced hepatitis, mice were i.p. injected with 25 µg/kg murine recombinant TNF (Peprotec) or 5 µg/kg LPS (Sigma) combined with 1000 mg/kg GalN (Sigma) in PBS for 6h (Hamesch et al., 2015). All mice were sacrificed by CO2 and cervical dislocation, liver lobes were stored in 10% formalin, embedded in cryo embedding matrix OCT (Roth), or shock-frozen in liquid nitrogen for further analysis. Serum from heart blood was obtained by coagulation and centrifugation for 15 min at 1000 x g. Serum Alanine aminotransferase (ALT) levels were measured using the Reflotron® Plus (Roche) with the respective ALT/GPT test stripes.

### Hepatocyte isolation

To obtain primary hepatocytes of LRH-1 L2L2 and AlbCre mice for *ex vivo* culture, the largest five liver lobes were isolated, short-term stored in 1:75 Heparin/PBS (Sigma) and perfused with 37 °C warm Buffer I (HBSS (Sigma), 0.1% Glucose and 2 mM EGTA) for 10 min. To digest the extracellular matrix, additional perfusion with Buffer II (HBSS (Sigma), 5 mM CaCl2 and 0.3 mg/ml Collagenase NB 4 Grade (Nordmark)) for 15-20 min and subsequently incubation for 5 min at 37°C was performed. The capsule of the lobes was disrupted using forceps in DMEM/F-12 Ham (Sigma) media with supplements (10% FCS (Sigma), 4 mM L-glutamine (Sigma-Aldrich), 1 x Pen/Strep 100x (Sigma-Aldrich), 50 µg/ml gentamycin (Sigma), 10 mg/ml Insulin Transferrin Sodium Selenite Supplement (ITS, Roche)) to receive a single cell suspension. After filtering through 100 µM strainer and centrifugation at 75 x g for 3 min, the hepatocyte pellet was resuspended in 40% Percoll (GE Healthcare)/medium and centrifuged at 500 x g for 10 min to remove dead cells. Live cells in the pellet were resuspended, counted, and seeded per well with 1.5 x 10^4^ cells per well for 96-well and Seahorse plate, 0.4 x 10^6^ cells/6-well and 1.5 x 10^6^ cells/10-cm dish. All plates were pre-coated with 0.3 mg/ml collagen (Sigma) for 2-24h prior to seeding. Hepatocytes were kept in culture at 37 °C with 5% CO2 with daily medium exchange and not longer than 3 days.

### LRH-1 overexpression cell line IHH

Immortalized human hepatocytes (IHH) cells (Nguyen et al., 2005) were cultured in same DMEM/F12 medium as hepatocytes. GEV16-mediated 4-hydroxytamoxiphen (4-HT)-inducible overexpression of human Flag-tagged LRH-1 was achieved as previously described (Vince et al., 2007). In brief, two-step cell line generation was performed by first transduction of target cells with HEK293T-dervided virus containing the GEV16 plasmid (pHCA/GAL4(1-93).ER.VP16) and generation of a stable cell line by selection with hygromycin B. Secondly, generated GEV16-cells were transduced with expression plasmids (pF-5xUAS-MCS-SV40-Puro-based) containing the human Flag-tagged LRH-1 gene, and selected with 1 µg/ml puromycin. Overexpression of target genes was achieved by treatment of cells with 4-HT 24h prior to experiments. Additional TNF treatment took place for 2 h.

### Histological analysis

Formalin-fixed and paraffin-embedded murine liver tissue was cut in 4 µm sections, rehydrated and stained by hematoxylin and eosin (H&E) with subsequent dehydration and mounting.

Connective tissue and collagen deposition in the hepatic tissue was visualized by Picro Sirius Red staining. Rehydrated paraffin sections were stained with Hematoxylin for 5 min and afterwards washed in running tap water for 5 min. Sections were incubated in Picro Sirius Red solution (0.1% Sirius Red (Sigma) in saturated, aqueous solution of picric acid) for 1h and washed twice in 0.5% acetic acid water before samples were dehydrated and mounted. Quantification of fibrosis was determined by measuring the percentage of red-stained area based on thresholds using ImageJ software.

Lipid deposition in hepatic tissue was determined by Oil Red O staining as described previously (Mehlem et al., 2013). Briefly, frozen OCT-embedded liver tissue was cut in 10 µm sections with Hydrax C500 cryostat (Zeiss), fixed in 10% formalin for 15 min and afterwards washed twice with water and with 60% isopropanol for 5 min. Sections were stained with filtered Oil Red O solution (15 ml of 0.3% OilRedO (Sigma) in isopropanol mixed with 10 ml H2O) for 15 min and washed with tab water until clear. Hematoxylin counterstaining was performed for 1 min and slides were mounted and sealed after washing. Quantification of lipids was determined by measuring the percentage of red-stained area based on thresholds using ImageJ software.

For cleaved Caspase 3 immunohistochemistry, antigens of paraffin-embedded and rehydrated liver sections were retrieved by boiling in 10 mM sodium citrate buffer (pH 6) for 15 min. Samples were further blocked in 3% hydrogen peroxide/PBS for 60 min, followed by incubation in 5% goat serum, 3% BSA in PBS for additional 60 min. Staining with 1:100 of rabbit anti-cleaved Caspase 3 (Cell Signaling Technologies) or goat anti-rabbit-IgG (Jackson Immuno) in second blocking solution took place over night at 4 °C, and was followed by 3 x 10 min wash in TBS-Tween and incubation with 1:100 biotinylated secondary goat anti-rabbit (Jackson Immuno) for 2 h, followed by additional 3 x 10 min wash and incubation in ABC solution of Vectastain ABC Kit (Vector laboratories). After washing, visualization was achieved by incubation with 1 x DAB substrate solution (Roche) for 5-10 min and washing in running tab water for 5 min. Counterstaining was performed in hematoxylin for 10 sec, followed by dehydration and mounting.

For RelA immunocytochemistry, isolated hepatocytes were seeded in glass-bottom µ-slides (IBIDI) and optionally treated with 20 ng/ml TNF for 1h to induce RelA nuclear translocation. Media was removed and cells were fixed in 10% formalin for 15 min, washed with PBS and permeabilized in 0.1% Triton X-100 for 15 min. Blocking of unspecific binding took place in blocking solution (3% BSA, 5% goat serum in 0.1% Saponin in PBS) for 60 min, whereafter cells were stained with 1:500 rabbit anti-RelA/p65 (Cell Signaling Technologies) in blocking solution overnight. After three times 5 min washing in PBS, cells were stained with 1:500 goat anti-rabbit Alexa Fluor 568 secondary antibody for 1.5 h. Following three times washing, cells were covered with Fluoroshield™ mounting medium containing DAPI (Sigma) to stain nuclei and mount samples.

### High-dimensional FACS analysis for immune cell phenotyping

To analyze hepatic immune cells, high-dimensional flow cytometry and unsupervised analysis with automated FlowSOM cluster identification was performed as described in Phan et al. (Phan et al., 2021). Non-parenchymal hepatic cells were isolated as described in Liu et al. (Liu et al., 2020). Liver lobes were isolated and incubated as small cut pieces in 3 ml Digestion Buffer (0.2 mg/ml Collagenase, 0.05 mg/ml DNase1, 10% FCS, DMEM media) for 1h at 37 °C in 6-well plates. Digested tissue was homogenized by pipetting and using a G20-syringe needle, afterwards it was filtered through 70 µm strainer with subsequent washing with cold DMEM media. Slow centrifugation at 50 x g for 3 min at 4 °C separated hepatocytes (pellet) and non-parenchymal cells (supernatant). After further centrifugation of supernatant at 350 x g for 7 min at 4 °C, erythrocyte lysis was performed by incubation in 3 ml ACK lysis buffer (0.15 M NH4Cl, 10 mM KHCO3, 0.1 mM Na2EDTA in H2O, pH 7.4) for 5 min. After centrifugation, pellet was washed twice with 10 ml cold PBS, resuspended, and cells were counted in 1 ml cold PBS. For myeloid lineage analysis, 12×10^6^ cells were stained with 1:1000 Fixable viability dye (VD455UV) (eBioscience) for 30 min at 4 °C, washed with FACS Buffer (2% BSA, 2 mM EDTA, 2 mM NaN3), blocked in 1:200 anti-CD16/32 TruStain Fcγ PLUS antibody (BioLegend) for 30 min at 4 °C, and stained with a combination of surface marker antibodies labelled with fluorophores (see Supplementary Table 1 and 2) for 45 min at 4 °C, washed with cold PBS and measured at LSR Fortessa (BD). For lymphoid lineage analysis, 1-2×10^6^ cells were stimulated for 5 h at 37 °C with 50 ng/ml PMA, 500 ng/ml Ionomycin and 10 µg/ml Brefeldin A, whereafter cells were stained with FVD455UV, blocked, and stained as described. For intracellular staining, cells were permeabilized and fixed with Foxp3/Transcription Factor Staining Buffer Set (eBioscience), intracellularly blocked as before, and stained with cytokine-specific antibodies labelled with fluorophores overnight at 4 °C. After washing, cells were measured at LSR Fortessa (BD). Compensation was calculated by FMO-controls and negative CompBeads (BD) controls in FlowJo software.

Compensated, cleaned, live, CD45+ subsets were exported from FlowJo and analyzed in R environment (version 4.1.1) using an adapted workflow based on described analyses (Brummelman et al., 2019; Ingelfinger et al., 2021; Nowicka et al., 2019). Briefly, FCS data were transformed by calculating with an inverse hyperbolic arcsinh function with individually set cofactors, normalized to values between 0 and 1, and then used to create Uniform Manifold Approximation and Projection (UMAP) plots according to umap package (version 0.2.8.0). Cell clusters were determined using FlowSOM (version 2.0.0) and ConsensusClusterPlus (version 1.56.0) and subsequently manually merged and attributed to the different hepatic immune cell population based on median marker expression.

### ELISA

To detect murine TNF levels, sandwich ELISA was performed following BioLegend’s instructions. Briefly, high affinity 96 well plates (Nunc) were coated with 3 µg/ml rat anti-TNF capture antibody (BioLegend) in coating buffer (0.1 M NaHCO3, 0.03 M Na2CO3, pH 9.5) overnight at 4 °C. After washing 4 times with 0.05% Tween/PBS, unspecific binding was blocked with 1% BSA/PBS for 2 h, whereafter 1:2-diluted samples and a recombinant protein standard (500 pg/ml to 0 pg/ml) were added and incubated overnight at 4 °C on a shaker. After washing, 1 µg/ml rat biotinylated anti-mouse TNF detection antibody (BioLegend) was added and incubated for 1h. After further washing, plates were incubated with streptavidin peroxidase-conjugate (Calbiochem) for 30 min, washed and then treated with trimethylboron substrate reagent (BioLegend) for signal development. Reaction was stopped using sulfuric acid and absorbance was measured at 450 nm with Infinite® 200 PRO plate reader (TECAN). Amount of TNF was calculated using the included TNF standard dilution series.

### RT-qPCR

RNA isolation of liver or cells was archived by lysing samples in peqGOLD TriFast™ (VWR) or TRIzol™ Reagent (Invitrogen) according to manufacturer’s protocol. Small pieces of frozen murine liver were lysed in 1 ml by 4 Hz shaking in the TissueLyzer II (QIAGEN). Harvested cell pellets were directly lysed in 1 ml peqGOLD TriFast™ (VWR) or 400 µl TRIzolTM Reagent (Invitrogen). After isolation of RNA, chromosomal DNA was digested with DNase I (NEB), and 2 µg of DNA-free RNA were reverse transcribed into cDNA by High-Capacity cDNA Reverse Transcription Kit (Applied Biosystems). Real-Time-quantitative PCR was performed on a StepOnePlus™ qPCR device (Applied Biosystems) using SYBR® Green (Applied Biosystems) and specific, exon-exon-spanning primers. Murine or human beta-actin CT values were used for normalization of respective samples (ΔCT(gene) = CT(gene) – CT(β-actin)).

### Western Blotting

For Western Blot analysis, small pieces of murine liver were homogenized in 1 ml RIPA Buffer (50 mM Tris pH 7.4, 150 mM NaCl, 0.1% SDS, 1% NP-40, 0.5% sodium deoxycholate, fresh 1x cOmplete™ Protease Inhibitor Cocktail (Roche)) by 4 Hz shaking in the TissueLyzer II (QIAGEN). Cultured hepatocytes were harvested from 6-well plates using Trypsin/EDTA (Sigma) and lysed in 100-200 µl RIPA Buffer on ice for 20 min. Lysates were centrifuged for 20 min at 14.000 x g, 4 °C and total protein concentration of supernatant was determined using Pierce™ BCA Assay Kit (Thermo Fisher). Samples for Western Blotting were boiled in 1x Laemmli buffer (5x: 5% SDS, 0.2 mM EDTA, 0.02% Bromophenol blue, 125 mM Tris pH 6.8, 50% Glycerol, 160 mM DTT) for 5 min at 95 °C and equal protein amounts (30-50 µg/sample) were loaded into 12% SDS-acrylamide gel pockets (stacking gel: 5% Rotiphorese® 30 Gel (Roth), 0.13 M Tris-HCl pH 6.8, 0.1% SDS, 0.1% APS, 0.01% TEMED; separation gel: 12% Rotiphorese® 30 Gel (Roth), 0.38 M Tris-HCl pH 8.8, 0.01% SDS, 0.01% APS, 0.08% TEMED). After SDS-Polyacrylamide-gel electrophoresis (SDS-PAGE), proteins were wet-blotted on a PVDF membrane (Roche) in Blotting Buffer (25 mM Tris, 192 mM glycine, 0.1% SDS, 21.5% methanol), unspecific binding was blocked in 5% non-fat dry milk/TBS-T (137 mM NaCl, 2.7 mM KCl, 15 mM Tris, 0.1% Tween-20) for 1h and the membrane was incubated with primary antibodies in 5% BSA/TBS-T over night at 4 °C. After 3 x 5 min wash intervals in TBS-T, incubation with peroxidase-conjugated secondary antibodies in 5% BSA/TBS-T took place for 2 h at room temperature with additional washing afterwards. For visualization, ECL solution (250 mM Luminol, 90 mM p-coumaric acid, 10 mM Tris) with 1% hydrogen peroxide was applied on membranes and proteins were detected in Image Quant LAS4000 (GE Heatlhcare).

### Cytosolic and mitochondrial fractionation

To study mitochondrial outer membrane permeabilization, mitochondrial and cytosolic fractions were collected by harvesting cells from 10-cm dishes using Trypsin/EDTA, lysing in 200 µl cold cytosolic extraction buffer (1.4 mM KH2PO4 ph7.2, 4.3 mM Na2HPO4, 250 mM sucrose, 70 mM KCl, 137 mM NaCl, 200 µg/ml digitonin, 1x cOmplete™ Protease Inhibitor Cocktail (Roche)) on ice for 5 min and collecting the supernatant (cytosolic fraction) after centrifugation for 10 min at 1000 x g, 4 °C. The pellet containing mitochondria was washed once with cytosolic extraction buffer by additional centrifugation and thereafter lysed in 200 µl mitochondrial lysis buffer (50 mM Tris pH 7.4, 150 mM NaCl, 2 mM EDTA, 2 mM EGTA, 0,2% Triton-X-100, 0,3% NP-40, 0,5% deoxycholic acid, 1x cOmplete™ Protease Inhibitor Cocktail (Roche)) on ice for 10 min. Mitochondrial faction was collected by centrifugation for 10 min at 14000 x g, 4 °C. Samples were boiled in 1 x Laemmli buffer as described and used in a 4:1 ratio (cytosolic/mitochondria) for Western Blotting.

### Immunoprecipitation

For immunoprecipitation (IP), cells were harvested from 10 cm dishes using Trypsin/EDTA, washed with PBS, lysed in 400 µl CHAPS buffer (50 mM Tris, 150 mM NaCl, 0,5% deoxycholic acid, 1% CHAPS, 1x cOmplete™ Protease Inhibitor Cocktail (Roche), pH 8.0) on ice for 30 min with subsequent centrifugation and protein determination as described in Western Blot method. As input control, 10% of supernatant was directly boiled in 1 x Laemmli buffer as described. Remaining supernatant was incubated with bait or isotype-antibodies for 2-3h at room temperature under constant rotation. Per sample 30 µl Protein G Sepharose 4 Fast Flow beads (GE Healthcare Life Sciences) were 3x times in CHAPS buffer by centrifugation at 14000 x g for 1 min each. Beads were added to samples and incubated overnight at 4 °C under constant rotation. On next day, beads were washed 3x to remove unbound proteins and boiled in 2x Laemmli buffer at 95°C for 10 min to release bound proteins. Equal volumes were used for Western Blotting.

### MTT cell viability assay

Cell death of hepatocytes was measured by 3-(4,5-dimethylthiazol-2-yl)-2,5-diphenyl-tetrazolium bromide (MTT, Sigma) viability assay. Hepatocytes were treated for 16h to induce cell death, whereafter medium was removed and cells were incubated in 10% MTT/media solution for 1h. Generated Violet formazan crystals of living cells were dissolved in DMSO, and absorbance was measured at 562 nm with the Infinite® 200 PRO plate reader (TECAN). Data was normalized to untreated cells of each genotype.

### Seahorse ATP Rate Assay

To quantify the cellular bioenergetic profile comprising glycolytic and mitochondrial ATP generation, Seahorse XF Real-Time ATP Rate Assay Kit (Agilent) was used following the manufacturer’s procedure. Briefly, hepatocytes were seeded into Seahorse XFe24 cell culture plates one day prior to measurement, medium was carefully replaced to Seahorse XF DMEM medium pH 7.4 (Agilent) supplemented with 18 mM glucose, 1 mM pyruvate and 2 mM L-glutamine, and cells were incubated at 37°C without CO2 for 60 min. Directly before measurement, medium was changed again. Seahorse extracellular Flux Analyzer XFe24 (Agilent) was started with pre-hydrated XFe24 sensor cartridge containing oligomycin in Port A (final 1.5 µM), a mix of rotenone, antimycin A (both final concentration 0.5 µM) and Hoechst33342 (final 1 µg/ml) in Port B. After measurement, the cell number was determined based on Hoechst33342-positive nuclei using an automated algorithm in Cellomics ArrayScan^TM^ VTI High-Content Screening (Cellomics^TM^). Nuclei were identified based on shape, intensity, and size. Data analysis was performed with Agilent Seahorse Analytics Online Software by normalizing to cell number and to 100% total ATP production (sum of mitoATP and gylcoATP) of untreated LRH-1 L2L2 hepatocytes.

### Lactate secretion

Lactate secretion was measured using the LAC 142 Kit (Diaglobal) and the Diaglobal Photometer the according to manufacturer’s protocol. For sampling, medium of hepatocytes, seeded in 6-well plates, was changed and supernatant samples were taken after 0, 2, 6 and 24h. Lactate secretion was calculated as slope over time normalized to LRH-1 L2L2 hepatocytes.

### Luciferase Assay reporter assay

To determine the transcription factor activity of LRH-1 and NF-kB in hepatocytes and IHH cells, luciferase reporter assays were performed as previously described with plasmids containing 5x LRH-1 response element (LRH-1 5xRE), 5x NF-kB RE or empty pGL3 vector as negative control (Schoonjans et al., 2002). Hepatocytes were seeded in 6-well plates, transiently transfected using Lipofectamine® 3000 transfection kit (Invitrogen) and optionally treated one day later with TNF for 2 h. Cells were lysed in lysis buffer (100 mM K2HPO4, 0.2% TritonX-100, 1 µM DTT, pH 7.8) for 15 min on ice, collected and centrifuged at 500 xg for 10 min. Supernatant was transferred in white 96-well plate and luminescence was assessed with Infinite® 200 PRO plate reader (TECAN) by automatically injecting ATP solution (10 mM ATP, 20 mM MgCl2, 35 mM Glycyl-gylcerine) and luciferin substrate (470 µM firefly luciferin, 270 µM coenzyme-A, 20 mM MgCl2, 35 mM Glycyl-gylcerine). IHH cells were seeded in 10 cm dishes and transfected with jetPEI (Polyplus) for 24h, then detached and distributed in 96-well plates, in which treatment with 4-Hydroxytamoxiphen (4-HT) for another 24h and afterwards with TNF for 2 h took place. Cell lysis and luciferase assay was performed as described. The co-transfected plasmid containing the β-galactosidase gene was utilized to enable normalization based on β-galactosidase-mediated color change reaction of o-nitrophenyl-b-D-galactopyranoside (ONPG) in Z-Buffer (0.2 mg/ml ONPG, 60 mM Na2HPO4*2H2O, 40 mM NaH2PO4*H2O, 10 mM KCl, 1 mM MgSO4*7H2O, 50 mM 2-mercaptoethanol). Luminescence was calculated as relative light units by normalizing light units obtained from luciferase assays with the absorbance at 405 nm obtained from the β-galactosidase assay.

### Protein cloning and purification for Microscale Thermophoresis

Full length human LRH1 (UniprotKB O00482) was cloned into a modified pETM-30 vector (EMBL collection) using MfeI and HindIII restriction sites. The pETM-30 vector was modified so as to replace the NcoI site located at bp 520 for an MfeI restriction site. The pETM-30 vector incorporates a His6-tag followed by a GST-tag and a TEV protease cleavage site N-terminal to the target construct. The ligand binding domain (LBD, residues 299-541) of human LRH-1 was cloned into the IPTG-inducible pETM-11 (EMBL collection) containing a N-terminal His6-tag followed by a tobacco etch virus (TEV) protease cleavage site. Full length human RelA (UniprotKB Q04206) was cloned into a modified pETM-41 vector (EMBL collection) using KpnI and MfeI restriction sites. The vector was modified to replace the NcoI site located at bp 4991 for an MfeI restriction site. The pETM-41 vector contains an His6-tag followed by an MBP-tag and a TEV protease cleavage site N-terminal to the target construct. The three different constructs of human RelA, namely RelA-R, RelA-RD, RelA-RDN, were obtained from the full-length RelA containing plasmid, that served as template, by applying site directed mutagenesis via whole plasmid PCR amplification using back-to-back primers. This procedure either removed unwanted domains or introduced stop codons to create the truncated constructs. All constructs were verified by sequencing (GATC, now Eurofins).

LRH1 and RelA constructs were expressed in E. coli Rosetta (DE3) cells (Millipore Sigma) in Luria Bertani medium supplemented with 25 μg/ml kanamycin and 35 μg/ml chloramphenicol. Cultures were grown at 37°C to an OD600 of 0.5. Protein expression was then induced by adding 0.25-0.5 mM isopropyl β-d-1-thiogalactopyranoside (IPTG) and growth continued at 18 °C for 16h. Cells were harvested by centrifugation at 5000 xg at 4°C for 20 minutes, pellets were resuspended in 30 ml lysis buffers (LRH1: 50 mM Tris pH 7.4, 500 mM NaCl, 5% glycerol, 1 mM DTT, 2 mM CAPS; LRH-1 LBD: 300 mM NaCl, 20 mM Tris-HCl pH 7.4, 1 mM CHAPS, 10% Glycerol, 5 mM DTT; RelA: 50 mM Tris pH 7.4, 300 mM NaCl, 1 mM DTT) supplemented with a EDTA-free protease inhibitor cocktail (Roche). Lysis was carried out by ultrasonication (Branson) for three times at 30-60 sec within 1.4 min pulse and 6/12 sec pulse at 40-50% transmission on ice. The homogenate was clarified by centrifugation at 40000 *g* for 90 min at 4 °C and the supernatant, supplemented with 20 mM imidazole, applied to a Ni^2+^-chelating His-Trap HP column (GE Healthcare) equilibrated in lysis buffer containing 20 mM imidazole. Samples were eluted using an increasing imidazole gradient (20 mM to 500 mM imidazole). LRH1 was then buffer exchanged into 50 mM Tris pH 7.4, 50 mM NaCl, 5% glycerol, 1 mM DTT, 2 mM CAPS before being applied to a Hi Trap Q HP (GE Healthcare) column and eluted with an increasing NaCl gradient. LRH-1 LBD and RelA constructs were dialyzed (buffer exchange) overnight at 4 °C in 0.5 L lysis buffer in the presence of TEV protease for His-tag removal. Afterwards constructs were subjected to subtractive metal affinity chromatography on a Ni^2+^-chelating His-Trap column (GE Healthcare). Fractions containing LRH-1 LBD were concentrated to 1.5 mL by centrifugal ultrafiltration using a 10 kDA Amicon Ultra tube (Millipore). The protein was further purified using a 16/60 Superdex 200 column (GE Healthcare) pre-equilibrated with GF Buffer (20 mM Tris-HCl pH 7.4, 150 mM NaCl, 1 mM CHAPS, 10% Glycerol, 10 mM DTT). RelA constructs were buffer exchanged into 50 mM Tris pH 7.4, 50 mM NaCl, 1 mM DTT and applied to a Hi-Trap SP HP (GE Healthcare) column and eluted with an increasing NaCl gradient. All protein samples were flash frozen in liquid nitrogen and stored at -80 °C until further use.

### Microscale Thermophoresis

Binding affinities between LRH-1 and RelA were assessed by microscale thermophoresis (MST) using the Monolith™ NT.115 instrument (NanoTemper Technologies). The purified LRH-1 (full length and LBD) were labeled with the Monolith NT™ Protein Labeling Kit GREEN-NHS according to manufactureŕs protocol. Before labeling, protein buffer was exchanged to 50 mM HEPES, pH 7.4 by gel filtration chromatography with a PD SpinTrap G-25 column (Cytiva). Unbound dye was removed and buffer was again exchanged to assay buffer (150 mM NaCl, 10 mM Tris-HCl (pH 7.4), 5% glycerol, 0.1% N-octyl β-D-glucopyranoside, 0.05% Tween-20) by overnight dialysis with a 3.5 kDa Slide-A-Lyzer MINI Dialysis Device (ThermoFisher Scientific). The labeled LRH-1 peptides were adjusted to 60 nM for the thermophoresis measurements. Amicon® Ultra Centrifugal Filter Devices (Millipore) were used to concentrate the purified RelA peptides to 80-85 µM and to exchange buffer to assay buffer. RelA peptides were 16-times 1:1 diluted with assay buffer with a final volume of 10 µl per dilution. Each dilution was mixed with 10 µl of labeled LRH-1 to gain a final LRH-1 concentration of 30 nM and final RelA concentration ranging from 40-42.5 µM to 1.2-1.3 nM. Samples were incubated for 30 min at room temperature in the dark and loaded into Monolith NT.115 capillaries (NanoTemper Technologies). MST was measured using a Monolith NT.115 instrument at 22°C, 60% LED power and 40% MST power. MST traces were analyzed with the KD binding model of MO Affinity Analysis software version 2.2.4 (NanoTemper Technologies). The cold region was defined as the 1 s region before the T-Jump while the hot region was defined as 10 s after the T-Jump for all samples.

### Statistics and data display

If not stated otherwise, data obtained from *in vivo* experiments are median ± standard deviation (SD) and data from *ex vivo* and *in vitro* experiments are means ± SD of a minimum of three independent experiments with several technical replicates. GraphPad Prism (version 8) software was used to perform statistical analysis and data visualization. Statistical significance was calculated by unpaired two-tailed Student’s test when comparing two groups, One-Way or Two-Way ANOVA when comparing three or more dependent groups, respectively. Suitable post-hoc tests (Tukey’s or Sidak’s) were applied and are stated in respective figure legends. Statistical significance is displayed as p-values with ≤ 0.05 being statistically significant (* = p < 0.05, ** = p < 0.01, *** = p < 0.001, **** = p < 0.0001).

---

**Figure S1.**
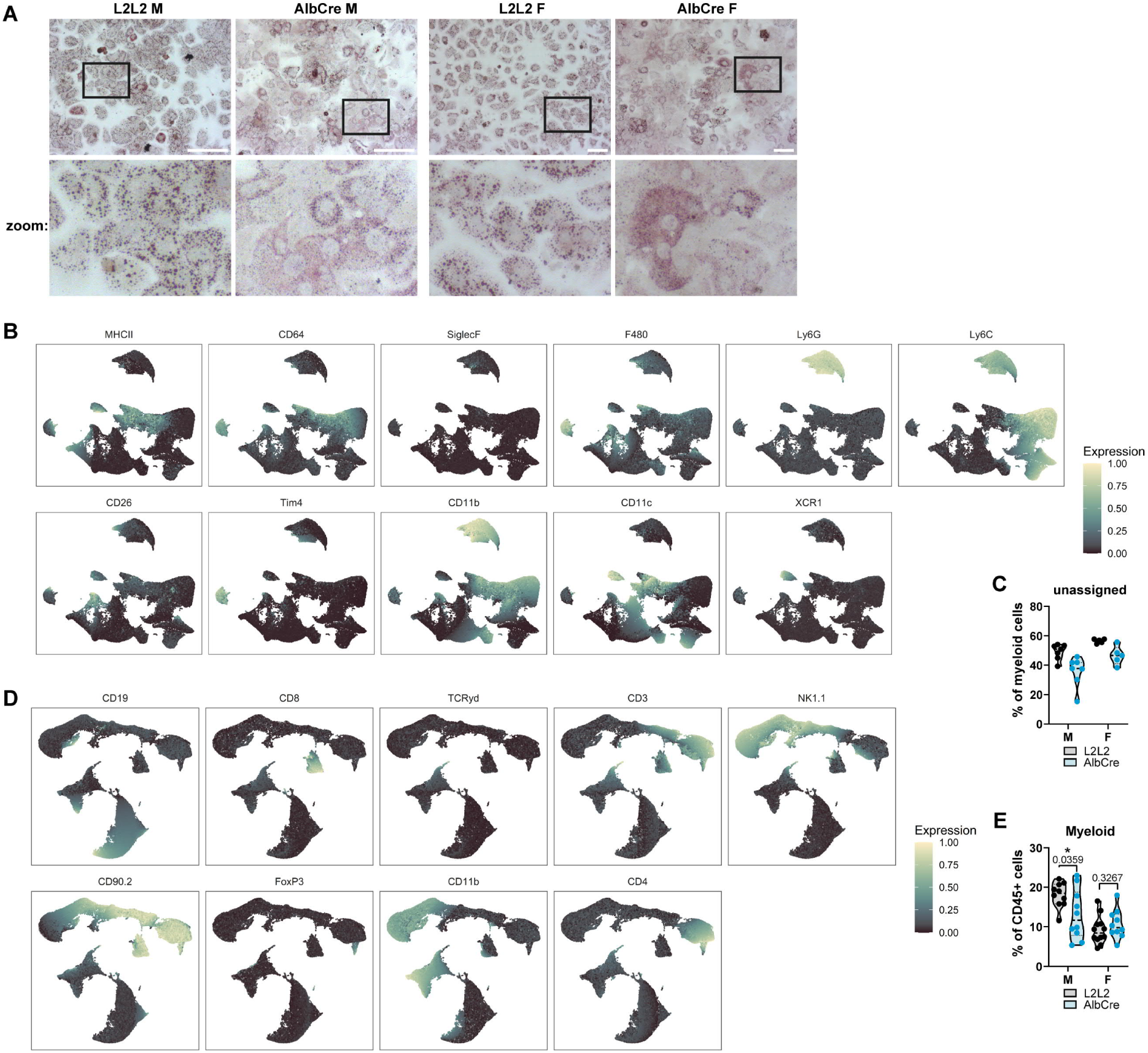
LRH-1 deficiency changes lipid distribution in hepatocytes. (A) Representative pictures of OilRedO-stained fat in hepatocytes of male (M) and female (F) LRH-1 L2L2 (L2L2) and LRH-1 AlbCre (AlbCre) mice. Scale bar 100 µm (left), 200 µm (right). (B-E) Computational flow cytometry analysis of liver non-parenchymal cells of mice described in (A). UMAP plots show single marker expression that were used for myeloid (B) and lymphoid (D) immune cell analysis. Percentage of undefined cells of each analysis are shown in (C) and (E). Violin blots show median and quartiles with each dot being one mouse. Statistical significance tested by Two-way ANOVA with Sidak’s multiple comparison test.

**Figure S2.**
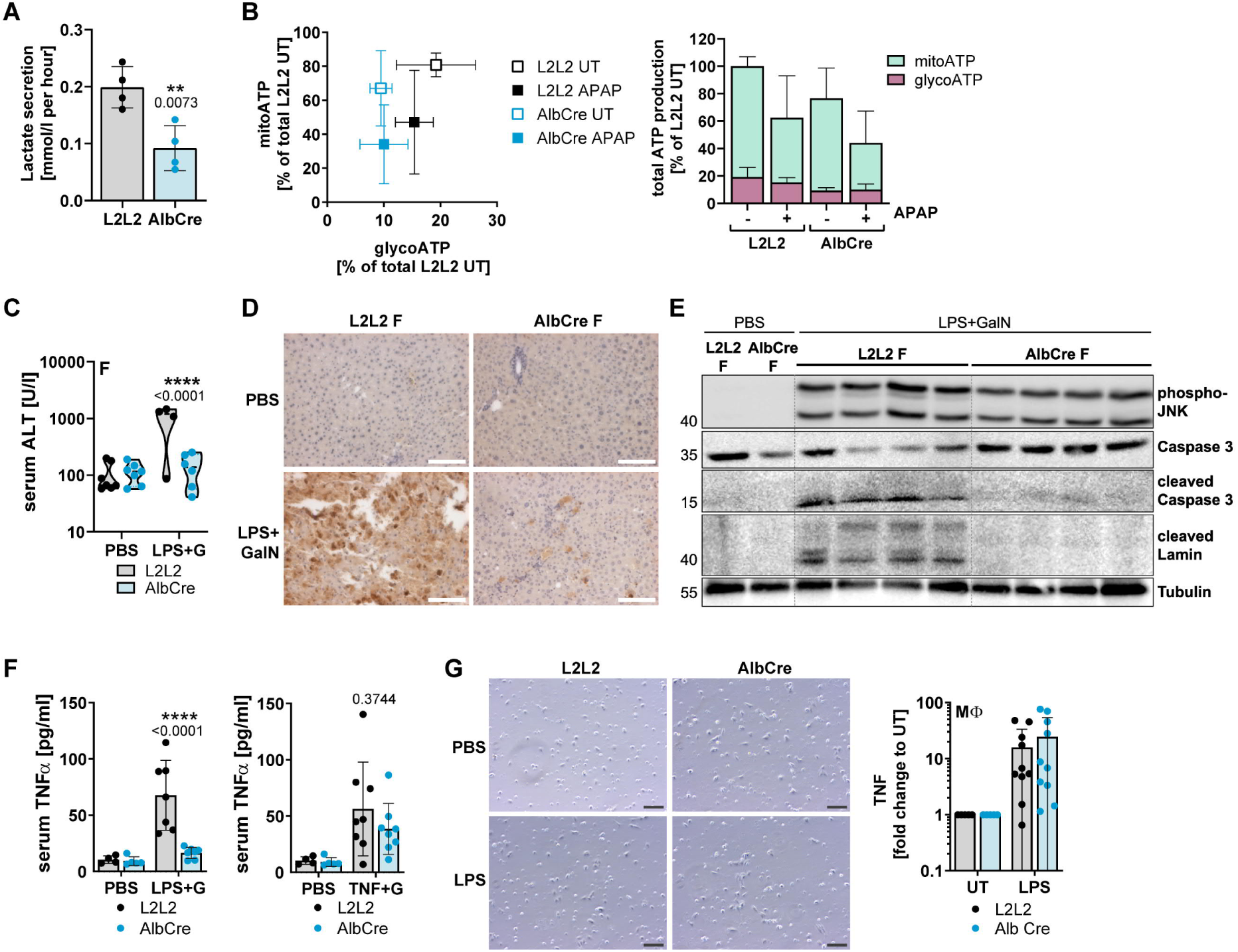
LRH-1 deficiency reduces ATP production and protects from LPS-induced liver damage. Data are from male (M) and female (F) LRH-1 L2L2 (L2L2) and LRH-1 AlbCre (AlbCre) mice. (A) Lactate assay of untreated hepatocytes. Lactate secretion was calculated as slope of time course of 2, 4, 6 and 24h. (B) Energy Map and bar graphs of Seahorse ATP Rate Assay performed in hepatocytes of LRH-1 L2L2 and AlbCre mice. APAP was injected by Seahorse device with final concentration of 15 mM. n=2. (C) Serum alanine aminotransferase (ALT) levels of mice treated with 5 µg/kg LPS and 1000 mg/kg N-Galactosamine (GalN, G) for 6h. (D) Cleaved Caspase 3 immunohistochemistry of liver section from mice treated as described in (C). Scale bar 250 µm. (E) Western Blot of liver lysates of mice treated as described in (C). Detected proteins are shown on the right, molecular weight is shown in kDa on the left. (F) Serum murine TNF levels determined by ELISA of mice treated as described in (LPS, C) or with 25 µg/kg TNF and 1000 mg/kg N-Galactosamine (TNF). (G) Picture of isolated hepatic macrophages and TNF levels in their supernatant measured by murine TNF ELISA. Macrophages were isolated, cultured for 24 day and then treated with 1 µg/ml LPS for 24h. Bar plots show mean +/- SD and violin blots show median and quartiles with each dot being one mouse. Statistical significance tested by Two-way ANOVA with Sidak’s multiple comparison test (C, F) or by unpaired student’s t test (A).

**Figure S3.**
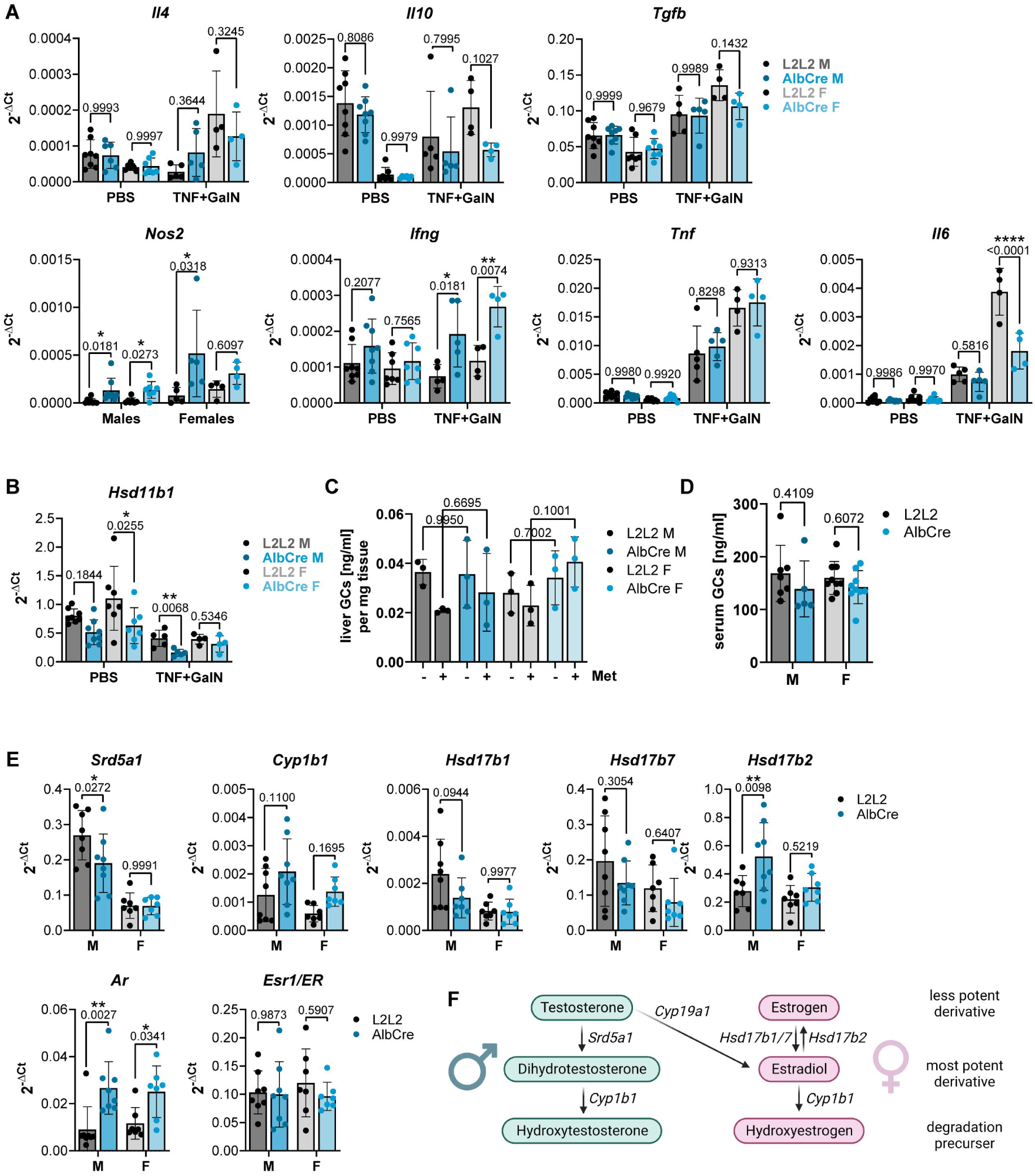
LRH-1 deficiency sex-dependently modulates several processes in the murine liver. Data are from male (M) and female (F) LRH-1 L2L2 (L2L2) and LRH-1 AlbCre (AlbCre) mice (A-B) Transcript levels of livers from mice treated with 25 µg/kg TNF and 1000 mg/kg N-Galactosamine (GalN). (C-D) Glucocorticoid (GCs) levels in ex vivo cultured liver (C) or in serum of untreated mice (D). Ex vivo cultured liver pieces were treated with 200 µg/ml Metyrapone (Met) to block *de novo* GCs synthesis and to show actual liver GCs levels. (E) Transcript levels of livers from untreated mice. (F) Scheme visualizing the role of sex hormone metabolizing enzymes that are shown in (E). Bar plots show mean +/- SD with each dot being one mouse. Statistical significance tested by Two-way ANOVA with Sidak’s multiple comparison test.

**Figure S4.**
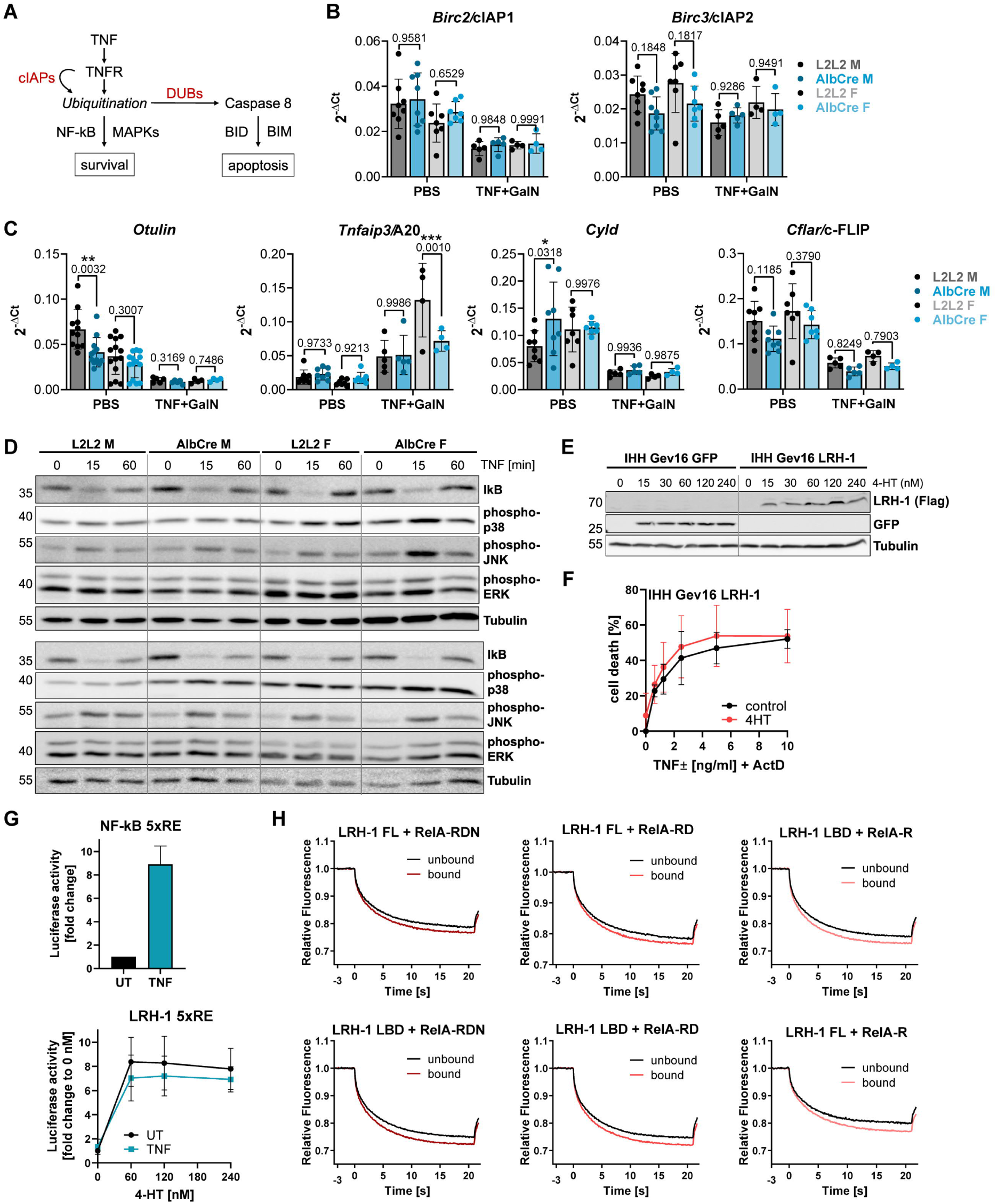
LRH-1 deficiency does not affect early TNFR signaling. Data are from male (M) and female (F) LRH-1 L2L2 (L2L2) and LRH-1 AlbCre (AlbCre) mice (A) Scheme visualizing TNF receptor (TNFR) pathway with ubiquitination of receptor complexes as critical decision point. cIAP1 (shown in (B)) promote ubiquitination and survival, while deubiquitinases (DUBs, shown in (C)) remove ubiquitination and promote apoptosis. (B-C) Transcript levels of livers from mice treated with 25 µg/kg TNF and 1000 mg/kg N-Galactosamine (GalN). (D) Western Blot of hepatocytes treated with 10 ng/ml murine TNF for indicated time. Detected proteins are shown on the right, molecular weight is shown in kDa on the left. n=2 (E) Western Blot of IHH Gev16 LRH-1 and GFP cells treated for 24h with 4-Hydroxytamoxifen (4-HT) to induce overexpression of human flag-tagged LRH-1 or GFP. Detected proteins are shown on the right, molecular weight is shown in kDa on the left. (F) MTT cell death assay of IHH Gev16 LRH-1 cell described in (E) treated for with 4-HT for 24h, and then with human TNF and 30 nM ActD for 24h. (G) LRH-1 and NF-kB activity measured by luciferase assay in IHH Gev16 LRH-1 or GFP cells described in (E). IHH cells were treated with 4-HT for 24h, then transfected with plasmids containing luciferase gene behind 5xRE of LRH-1 (low) or NF-kB (up) for 24h, then treated with 10 ng/ml human TNF for 1h and analyzed after 24h. n=6. (H) Microscale Thermophoresis fluorescence patterns visualizing dissociation properties of fluorescence-labelled full length LRH-1 or LRH-1 Ligand binding domain (LBD) peptides in combination with either RelA-RDN, RelA-RD or RelA-R peptides (explained in Main Figure 4E). Bar plots and line graphs show mean +/- SD with each dot being one mouse or one independent experiment. Statistical significance tested by Two-way ANOVA with Sidak’s multiple comparison test.

**Figure S5.**
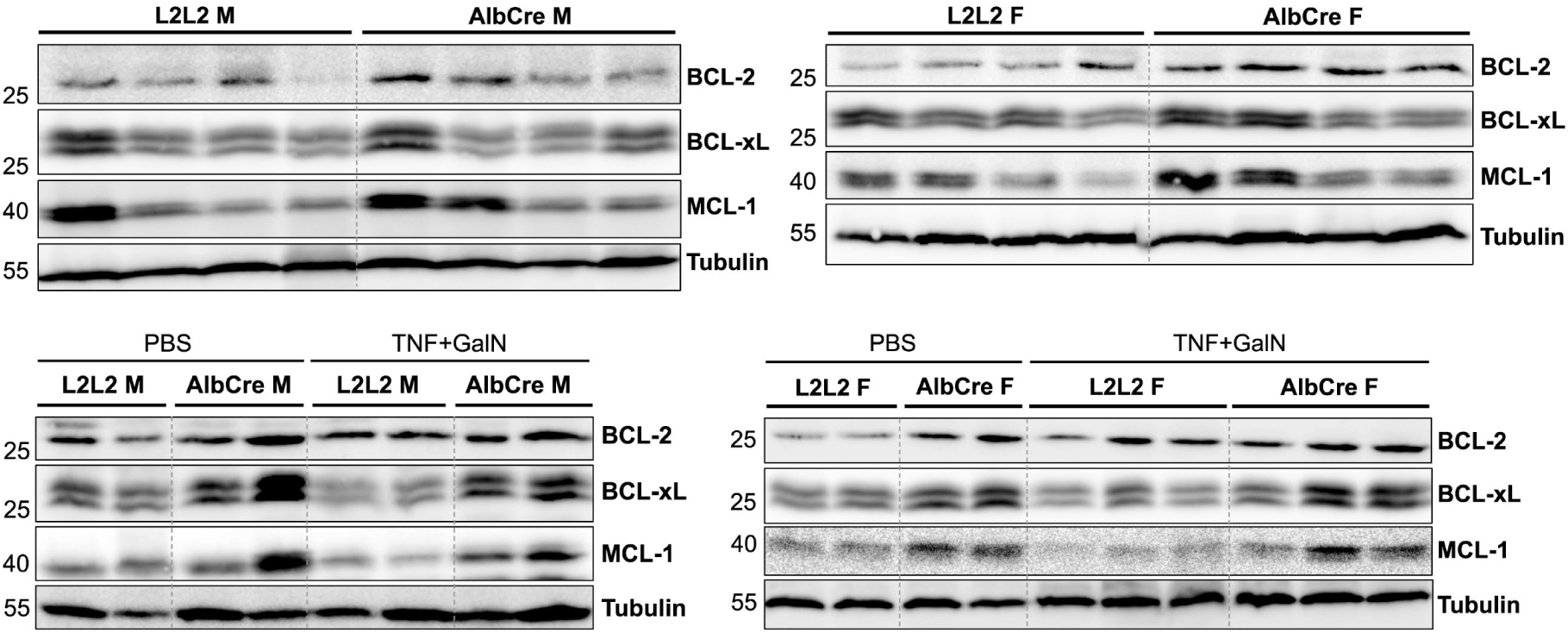
LRH-1 deficiency leads to upregulation of anti-apoptotic BCL-2 homologs. Western blot of liver from male (M) and female (F) LRH-1 L2L2 (L2L2) and LRH-1 AlbCre (AlbCre) mice untreated (upper panel) or treated with 25 µg/kg TNF and 1000 mg/kg N-Galactosamine (GalN) (lower panel) Only untreated samples were considered for quantification of protein expression shown in main Figure 5. Detected proteins are shown on the right, molecular weight is shown in kDa on the left.

**Figure S6.**
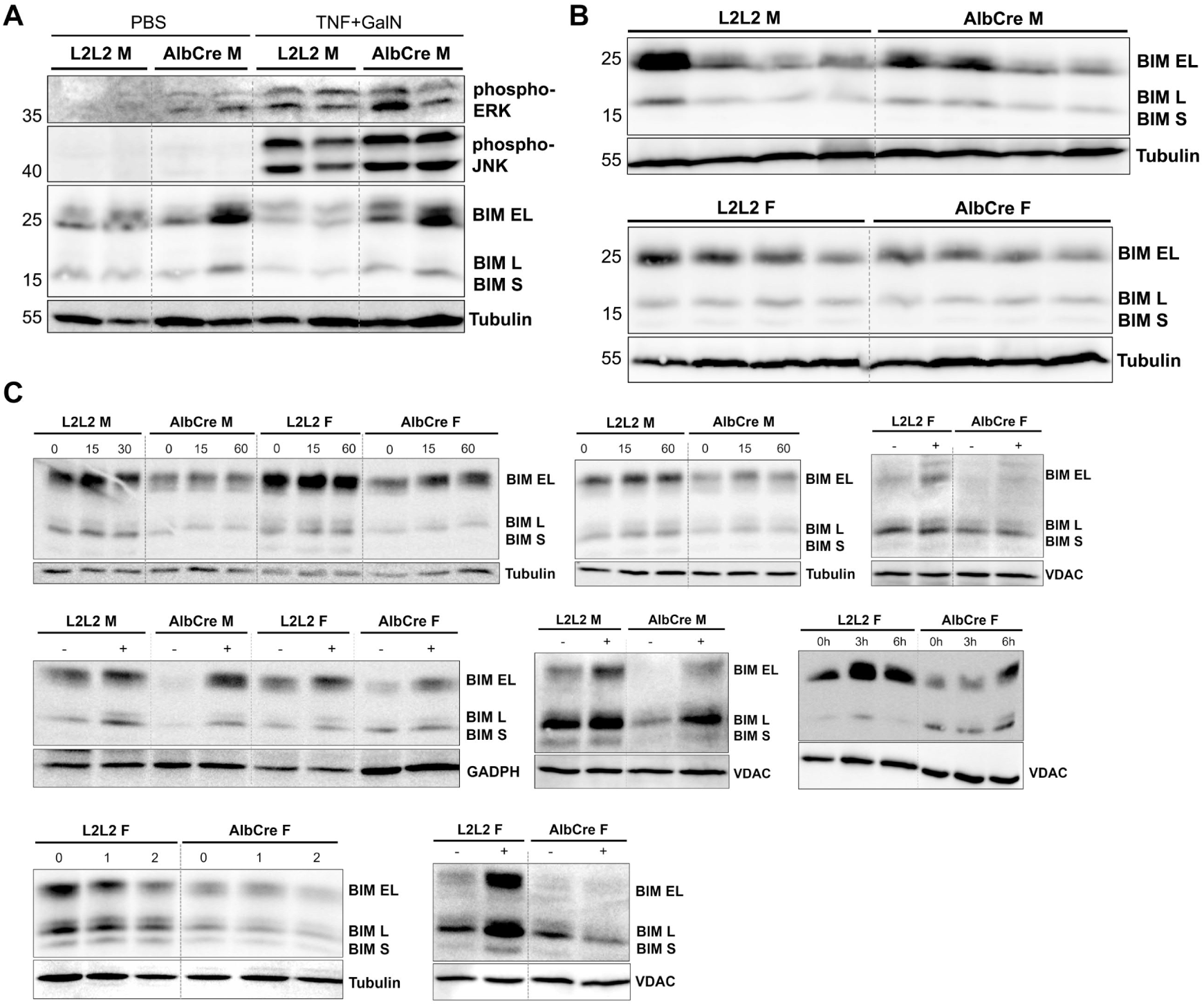
LRH-1 deficiency leads to downregulation of pro-apoptotic BCL-2 protein BIM. (A-B) Western Blots of liver from male (M) and female (F) LRH-1 L2L2 (L2L2) and LRH-1 AlbCre (AlbCre) mice treated with 25 µg/kg TNF and 1000 mg/kg N-Galactosamine (GalN) (A) or from untreated mice (B). (C) Western Blots of isolated hepatocytes from untreated mice described in (A-B). For all conditions, only untreated samples were considered for quantification of protein expression shown in main Figure 6. Detected proteins are shown on the right, molecular weight is shown in kDa on the left.

## Supplementary Tables

**Table 1:**
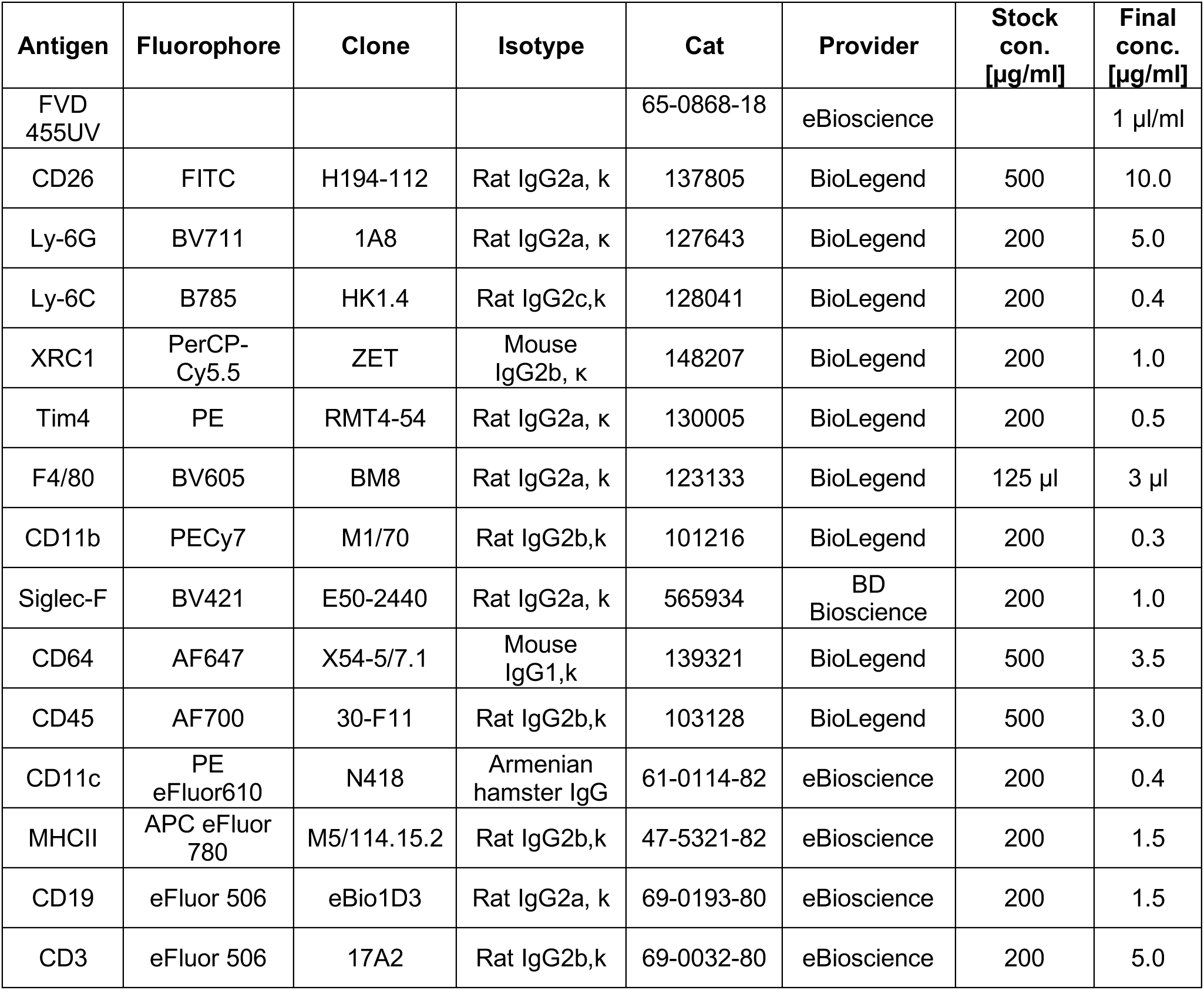
Antibodies used for myeloid panel in high dimensional FACS analysis.

| Antigen | Fluorophore | Clone | Isotype | Cat | Provider | Stock con.<br>[µg/ml] | Final conc.<br>[µg/ml] |
| --- | --- | --- | --- | --- | --- | --- | --- |
| FVD 455UV |  |  |  | 65-0868-18 | eBioscience |  | 1 µl/ml |
| CD26 | FITC | H194-112 | Rat IgG2a, κ | 137805 | BioLegend | 500 | 10.0 |
| Ly-6G | BV711 | 1A8 | Rat IgG2a, κ | 127643 | BioLegend | 200 | 5.0 |
| Ly-6C | B785 | HK1.4 | Rat IgG2c,κ | 128041 | BioLegend | 200 | 0.4 |
| XRC1 | PerCP-Cy5.5 | ZET | Mouse IgG2b, κ | 148207 | BioLegend | 200 | 1.0 |
| Tim4 | PE | RMT4-54 | Rat IgG2a, κ | 130005 | BioLegend | 200 | 0.5 |
| F4/80 | BV605 | BM8 | Rat IgG2a, κ | 123133 | BioLegend | 125 µl | 3 µl |
| CD11b | PECy7 | M1/70 | Rat IgG2b,κ | 101216 | BioLegend | 200 | 0.3 |
| Siglec-F | BV421 | E50-2440 | Rat IgG2a, κ | 565934 | BD Bioscience | 200 | 1.0 |
| CD64 | AF647 | X54-5/7.1 | Mouse IgG1,κ | 139321 | BioLegend | 500 | 3.5 |
| CD45 | AF700 | 30-F11 | Rat IgG2b,κ | 103128 | BioLegend | 500 | 3.0 |
| CD11c | PE eFluor610 | N418 | Armenian hamster IgG | 61-0114-82 | eBioscience | 200 | 0.4 |
| MHCII | APC eFluor 780 | M5/114.15.2 | Rat IgG2b,κ | 47-5321-82 | eBioscience | 200 | 1.5 |
| CD19 | eFluor 506 | eBio1D3 | Rat IgG2a, κ | 69-0193-80 | eBioscience | 200 | 1.5 |
| CD3 | eFluor 506 | 17A2 | Rat IgG2b,κ | 69-0032-80 | eBioscience | 200 | 5.0 |

**Table 2:** Antibodies used for lymphoid panel in high dimensional FACS analysis.

| **Antigen** | **Fluorophore** | **Clone** | **Isotype** | **Cat** | **Provider** | **Stock con.**<br>**[µg/ml]** | **Final conc.**<br>**[µg/ml]** |
| --- | --- | --- | --- | --- | --- | --- | --- |
| FVD 455UV |  |  |  | 65-0868-18 | eBioscience |  | 1 µl/ml |
| CD45 | FITC | 30-F11 | Rat IgG2b, k | 11-0451-85 | eBioscience | 500 | 3.3 |
| CD11b | PECy7 | M1/70 | Rat IgG2b,k | 101216 | BioLegend | 200 | 0.3 |
| CD3 | BV605 | 17A2 | Rat IgG2b, k | 100237 | BioLegend | 100 | 2.5 |
| CD19 | APC-Fire750 | 6D5 | Rat IgG2a, κ | 115557 | BioLegend | 200 | 2.0 |
| CD4 | PE-Dazzle594 | GK1.5 | Rat IgG2b, κ | 100455 | BioLegend | 200 | 0.5 |
| CD8 | BV421 | 53-6.7 | Rat IgG2a, k | 100737 | BioLegend | 125 µl | 1.3 |
| NK1.1 | BV711 | PK136 | Mouse IgG2a,k | 108745 | BioLegend | 200 | 1.6 |
| CD90.2 | BV786 | 30-H12 | Rat IgG2b, κ | 105331 | BioLegend | 200 | 3.0 |
| TCRyd | BV510 | GL3 | Armenian Hamster | 118131 | BioLegend | 500 | 2.5 |
| FoxP3 | PE | FJK-16s | Rat IgG2a, k | 12-5773-80 | eBioscience | 500 | 5 |
| IFNγ | APC | XMG1.2 | Rat IgG1, κ | 505809 | BioLegend | 200 | 5.0 |
| IL-17a | PerCP-Cy5.5 | eBio17B7 | Rat IgG2a, k | 45-7177-82 | eBioscience | 200 | 0.6 |

## KEY RESOURCES TABLE

The table highlights the reagents, genetically modified organisms and strains, cell lines, software, instrumentation, and source data **essential** to reproduce results presented in the manuscript. Depending on the nature of the study, this may include standard laboratory materials (i.e., food chow for metabolism studies, support material for catalysis studies), but the table is **not** meant to be a comprehensive list of all materials and resources used (e.g., essential chemicals such as standard solvents, SDS, sucrose, or standard culture media do not need to be listed in the table). **Items in the table must also be reported in the method details section within the context of their use.** To maximize readability, the number of **oligonucleotides and RNA sequences** that may be listed in the table is restricted to no more than 10 each. If there are more than 10 oligonucleotides or RNA sequences to report, please provide this information as a supplementary document and reference the file (e.g., See Table S1 for XX) in the key resources table.

***Please note that ALL references cited in the key resources table must be included in the references list.*** Please report the information as follows:

- **REAGENT or RESOURCE:** Provide full descriptive name of the item so that it can be identified and linked with its description in the manuscript (e.g., provide version number for software, host source for antibody, strain name). In the experimental models section (applicable only to experimental life science studies), please include all models used in the paper and describe each line/strain as: model organism: name used for strain/line in paper: genotype. (i.e., Mouse: OXTR^fl/fl^: B6.129(SJL)-Oxtr^tm1.1Wsy/J^). In the biological samples section (applicable only to experimental life science studies), please list all samples obtained from commercial sources or biological repositories. Please note that software mentioned in the methods details or data and code availability section needs to also be included in the table. See the sample tables at the end of this document for examples of how to report reagents.
- **SOURCE:** Report the company, manufacturer, or individual that provided the item or where the item can be obtained (e.g., stock center or repository). For materials distributed by Addgene, please cite the article describing the plasmid and include “Addgene” as part of the identifier. If an item is from another lab, please include the name of the principal investigator and a citation if it has been previously published. If the material is being reported for the first time in the current paper, please indicate as “this paper.” For software, please provide the company name if it is commercially available or cite the paper in which it has been initially described.
- **IDENTIFIER:** Include catalog numbers (entered in the column as “Cat#” followed by the number, e.g., Cat#3879S). Where available, please include unique entities such as <u>RRIDs</u>, Model Organism Database numbers, accession numbers, and PDB, CAS, or CCDC IDs. For antibodies, if applicable and available, please also include the lot number or clone identity. For software or data resources, please include the URL where the resource can be downloaded. Please ensure accuracy of the identifiers, as they are essential for generation of hyperlinks to external sources when available. Please see the Elsevier <u>list of data repositories</u> with automated bidirectional linking for details. When listing more than one identifier for the same item, use semicolons to separate them (e.g., Cat#3879S; RRID: AB_2255011). If an identifier is not available, please enter “N/A” in the column.

⚬ ***A NOTE ABOUT RRIDs:*** We highly recommend using RRIDs as the identifier (in particular for antibodies and organisms but also for software tools and databases). For more details on how to obtain or generate an RRID for existing or newly generated resources, please <u>visit the RII</u> or <u>search for RRIDs</u>.

Please use the empty table that follows to organize the information in the sections defined by the subheading, skipping sections not relevant to your study. Please do not add subheadings. To add a row, place the cursor at the end of the row above where you would like to add the row, just outside the right border of the table. Then press the ENTER key to add the row. Please delete empty rows. Each entry must be on a separate row; do not list multiple items in a single table cell. Please see the sample tables at the end of this document for relevant examples in the life and physical sciences of how reagents and instrumentation should be cited.

## TABLE FOR AUTHOR TO COMPLETE

*Please upload the completed table as a separate document. **<u>Please do not add subheadings to the key resources</u> <u>table.</u>** If you wish to make an entry that does not fall into one of the subheadings below, please contact your handling editor. **<u>Any subheadings not relevant to your study can be skipped.</u>** (**NOTE:** For authors publishing in Cell Genomics, Cell Reports Medicine, Current Biology, and Med, please note that references within the KRT should be in numbered style rather than Harvard.)*

## Key resources table

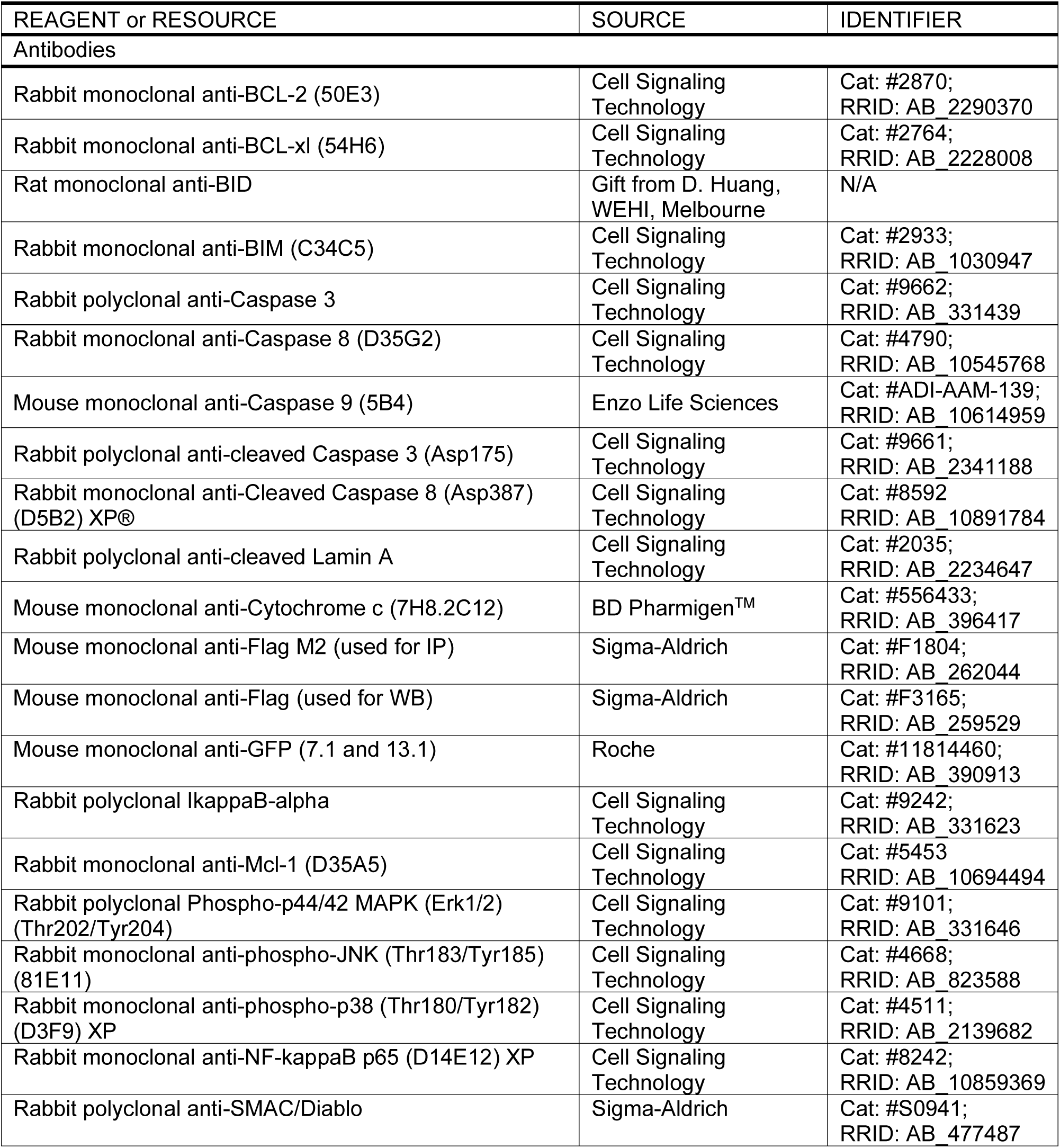
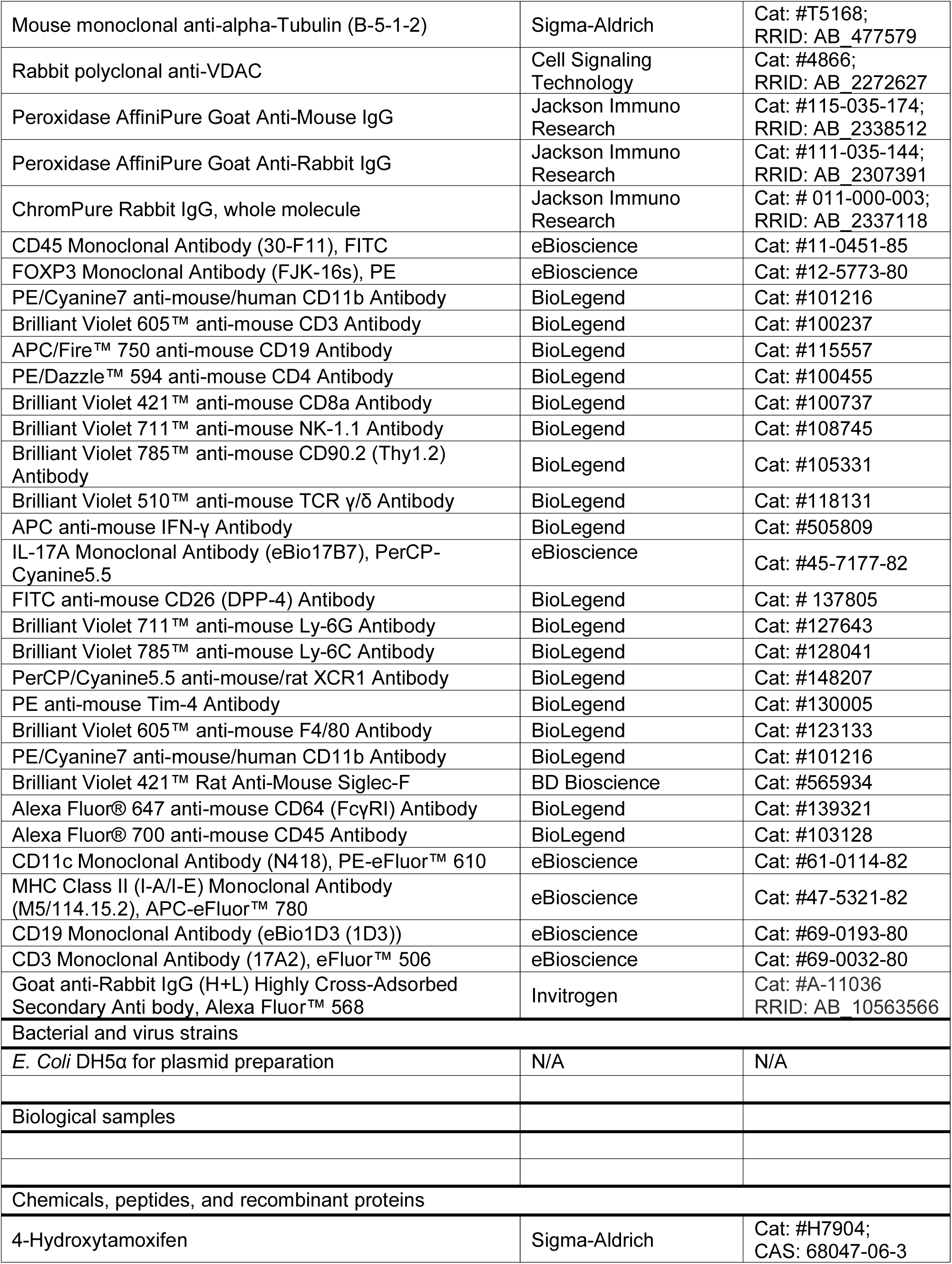
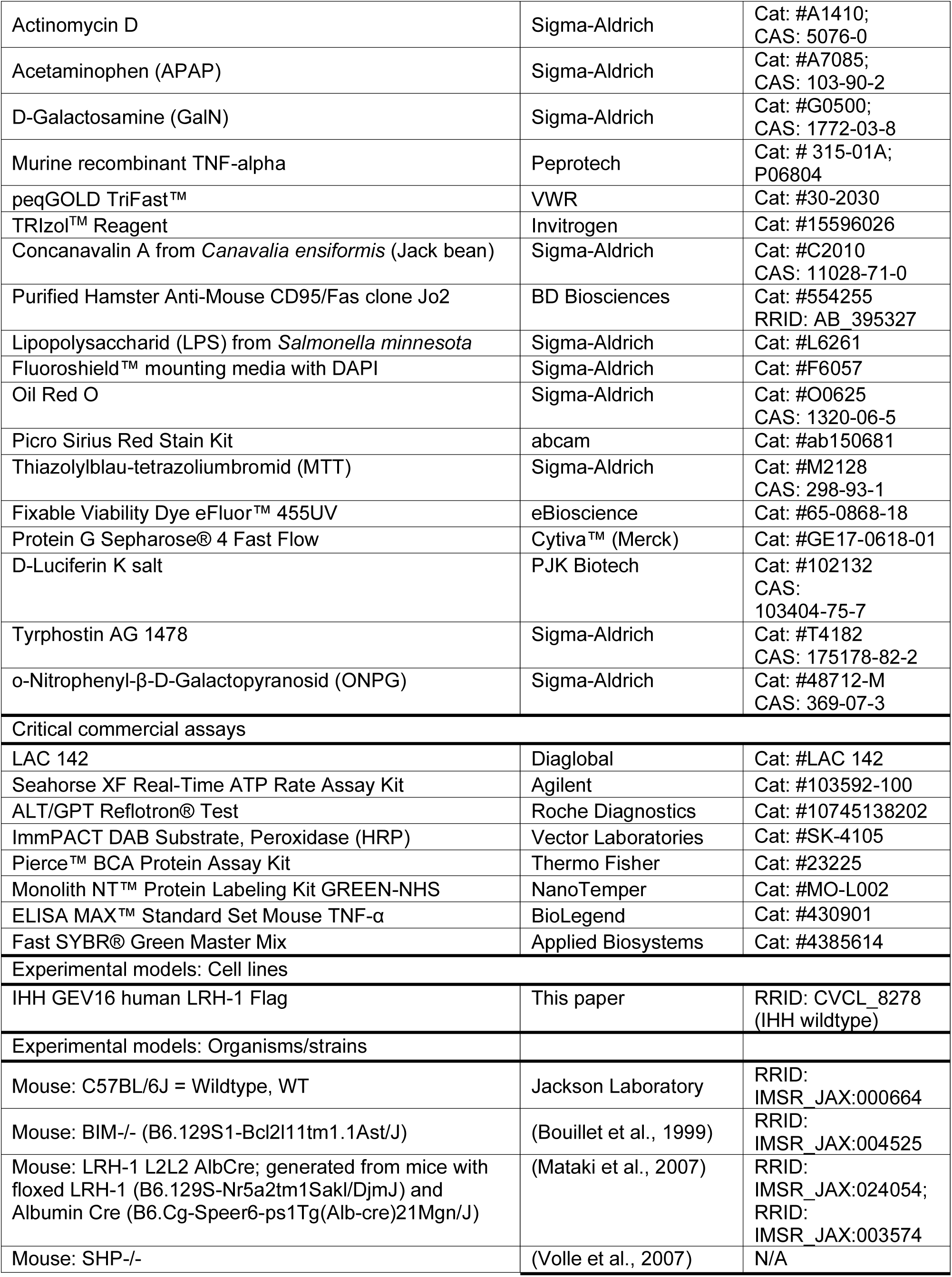
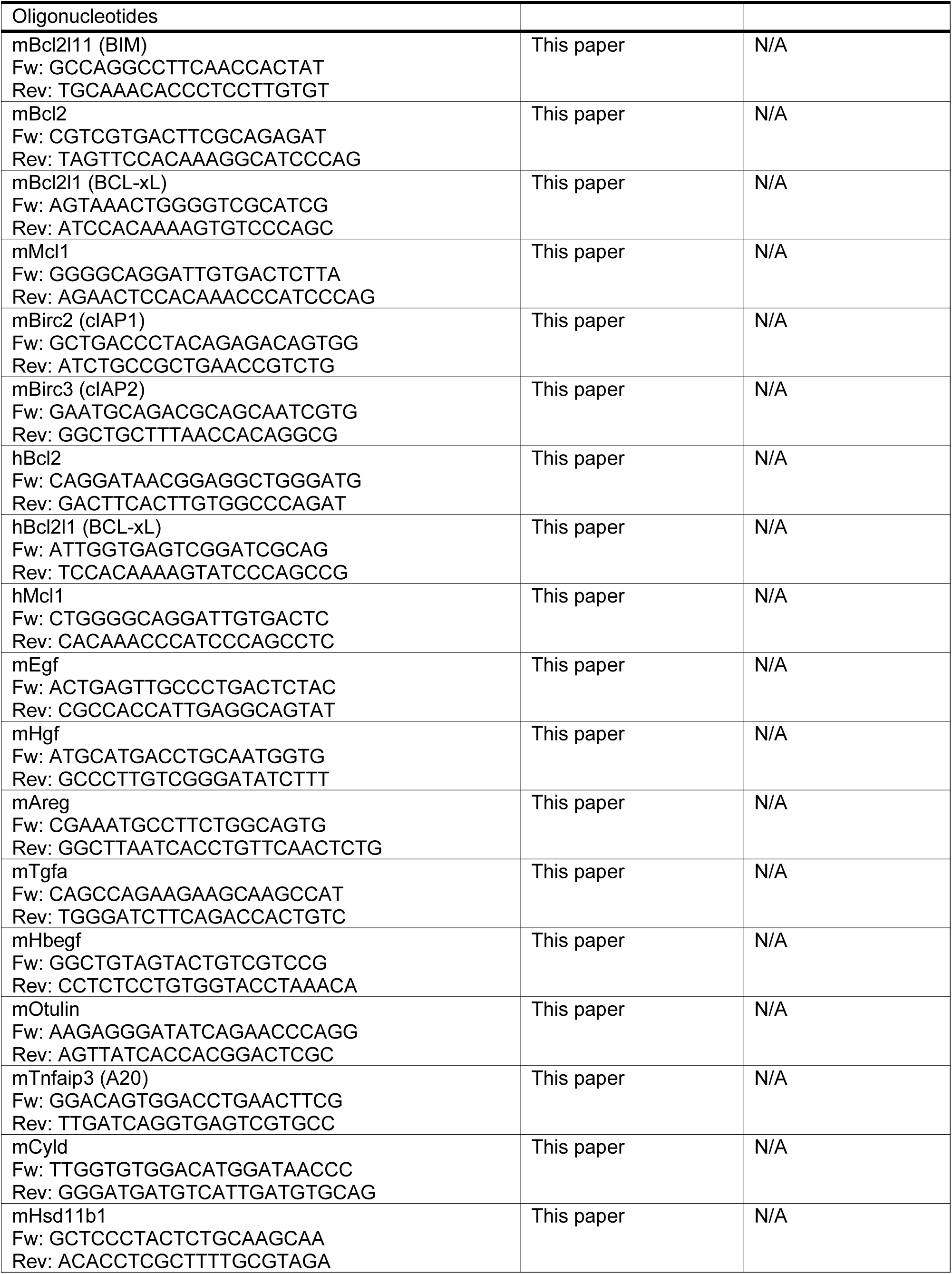
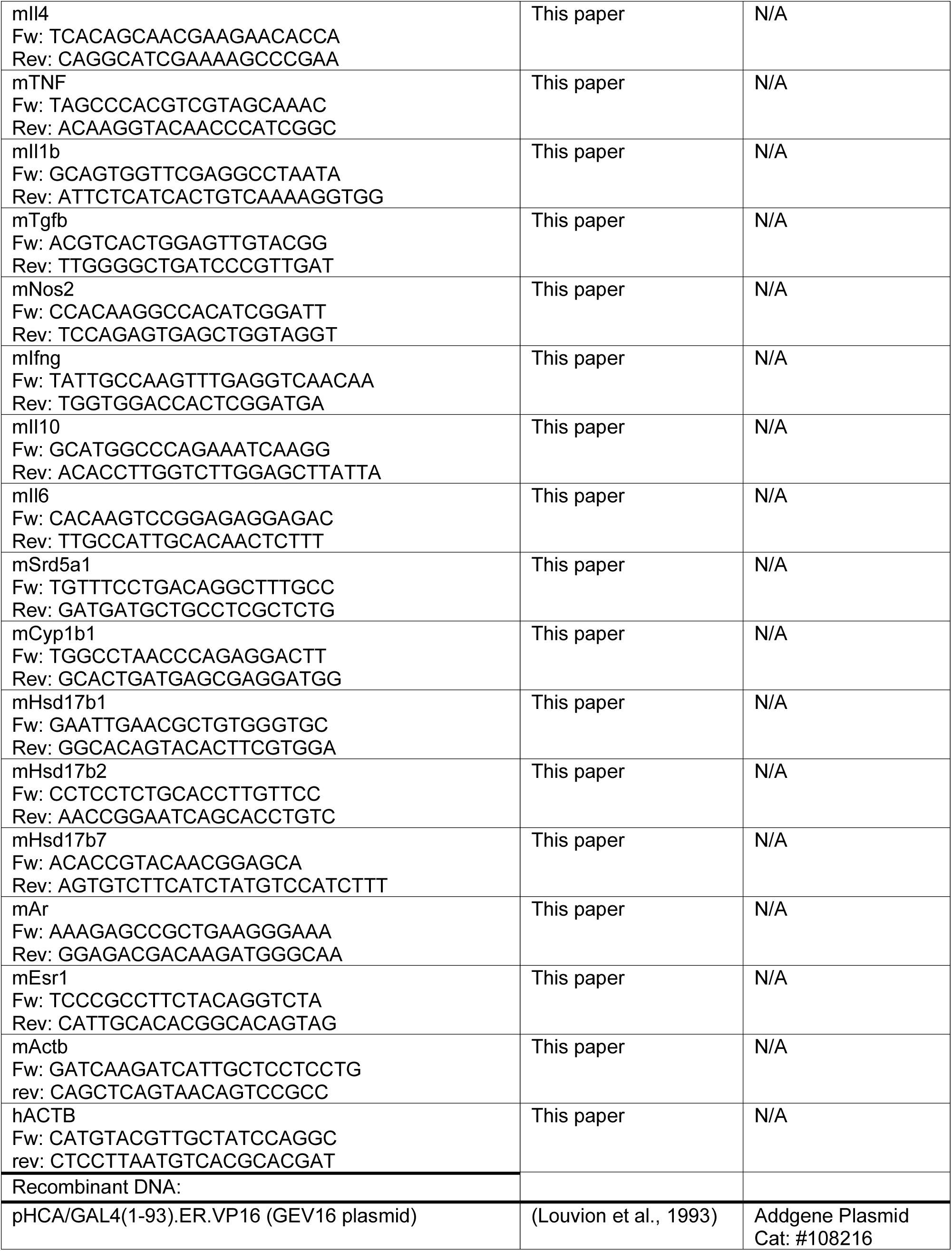
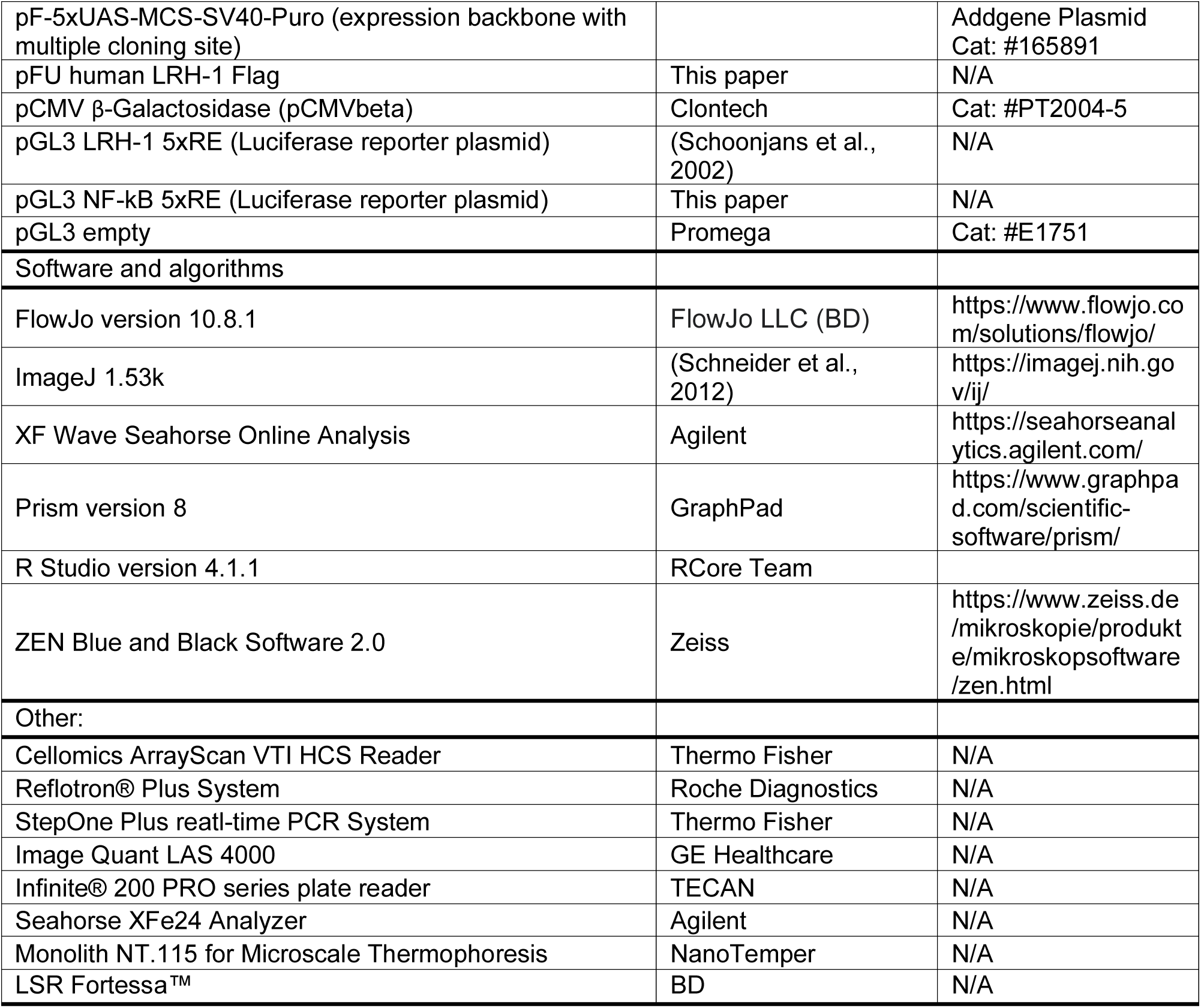

## References

Annicotte, J.-S., Chavey, C., Servant, N., Teyssier, J., Bardin, A., Licznar, A., Badia, E., Pujol, P., Vignon, F., Maudelonde, T., et al. (2005). The nuclear receptor liver receptor homolog-1 is an estrogen receptor target gene. Oncogene 24, 8167–8175. https://doi.org/10.1038/sj.onc.1208950.

Badmann, A., Keough, A., Kaufmann, T., Bouillet, P., Brunner, T., and Corazza, N. (2011). Role of TRAIL and the pro-apoptotic Bcl-2 homolog Bim in acetaminophen-induced liver damage. Cell Death Dis. 2, e171–e171. https://doi.org/10.1038/cddis.2011.55.

Bayrer, J.R., Wang, H., Nattiv, R., Suzawa, M., Escusa, H.S., Fletterick, R.J., Klein, O.D., Moore, D.D., and Ingraham, H.A. (2018). LRH-1 mitigates intestinal inflammatory disease by maintaining epithelial homeostasis and cell survival. Nat. Commun. 9, 4055. https://doi.org/10.1038/s41467-018-06137-w.

Ben-Moshe, S., Veg, T., Manco, R., Dan, S., Papinutti, D., Lifshitz, A., Kolodziejczyk, A.A., Bahar Halpern, K., Elinav, E., and Itzkovitz, S. (2022). The spatiotemporal program of zonal liver regeneration following acute injury. Cell Stem Cell 29, 973–989.e10. https://doi.org/10.1016/j.stem.2022.04.008.

Botrugno, O.A., Fayard, E., Annicotte, J.-S., Haby, C., Brennan, T., Wendling, O., Tanaka, T., Kodama, T., Thomas, W., Auwerx, J., et al. (2004). Synergy between LRH-1 and β-Catenin Induces G1 Cyclin-Mediated Cell Proliferation. Mol. Cell 15, 499–509. https://doi.org/10.1016/j.molcel.2004.07.009.

Bouillet, P., Metcalf, D., and Huang, D.C.S. (1999). Proapoptotic Bcl-2 relative Bim required for certain apoptotic responses, leukocyte homeostasis, and to preclude autoimmunity. Science (80-.). 286, 1735–1738. https://doi.org/10.1126/science.286.5445.1735.

Brewster, R.C., Weinert, F.M., Garcia, H.G., Song, D., Rydenfelt, M., and Phillips, R. (2014). The Transcription Factor Titration Effect Dictates Level of Gene Expression. Cell 156, 1312–1323. https://doi.org/10.1016/j.cell.2014.02.022.

Brummelman, J., Haftmann, C., Núñez, N.G., Alvisi, G., Mazza, E.M.C., Becher, B., and Lugli, E. (2019). Development, application and computational analysis of high-dimensional fluorescent antibody panels for single-cell flow cytometry. Nat. Protoc. 14, 1946–1969. https://doi.org/10.1038/s41596-019-0166-2.

Campana, L., Esser, H., Huch, M., and Forbes, S. (2021). Liver regeneration and inflammation: from fundamental science to clinical applications. Nat. Rev. Mol. Cell Biol. 22, 608–624. https://doi.org/10.1038/s41580-021-00373-7.

Chand, A.L., Wijayakumara, D.D., Knower, K.C., Herridge, K.A., Howard, T.L., Lazarus, K.A., and Clyne, C.D. (2012). The Orphan Nuclear Receptor LRH-1 and ERα Activate GREB1 Expression to Induce Breast Cancer Cell Proliferation. PLoS One 7, e31593. https://doi.org/10.1371/journal.pone.0031593.

Chen, C., Edelstein, L.C., and Gélinas, C. (2000). The Rel/NF-κB Family Directly Activates Expression of the Apoptosis Inhibitor Bcl-x L. Mol. Cell. Biol. 20, 2687–2695. https://doi.org/10.1128/MCB.20.8.2687-2695.2000.

Chong, H.K., Biesinger, J., Seo, Y.-K., Xie, X., and Osborne, T.F. (2012). Genome-wide analysis of hepatic LRH-1 reveals a promoter binding preference and suggests a role in regulating genes of lipid metabolism in concert with FXR. BMC Genomics 13, 51. https://doi.org/10.1186/1471-2164-13-51.

Cobo-Vuilleumier, N., Lorenzo, P.I., Rodríguez, N.G., Herrera Gómez, I. de G., Fuente-Martin, E., López-Noriega, L., Mellado-Gil, J.M., Romero-Zerbo, S.-Y., Baquié, M., Lachaud, C.C., et al. (2018a). LRH-1 agonism favours an immune-islet dialogue which protects against diabetes mellitus. Nat.Commun. 9, 1488. https://doi.org/10.1038/s41467-018-03943-0.

Cobo-Vuilleumier, N., Lorenzo, P.I., García Rodríguez, N., De, I., Herrera Gómez, G., Fuente-Martin, E., López-Noriega, L., Mellado-Gil, J.M., Romero-Zerbo, S.-Y., Baquié, M., et al. (2018b). LRH-1 agonism favours an immune-islet dialogue which protects against diabetes mellitus. Nat. Commun. 9, 1488. https://doi.org/10.1038/s41467-018-03943-0.

Cockram, P.E., Kist, M., Prakash, S., Chen, S.-H., Wertz, I.E., and Vucic, D. (2021). Ubiquitination in the regulation of inflammatory cell death and cancer. Cell Death Differ. 28, 591–605. https://doi.org/10.1038/s41418-020-00708-5.

Corazza, N., Jakob, S., Schaer, C., Frese, S., Keogh, A., Stroka, D., Kassahn, D., Torgler, R., Mueller, C., Schneider, P., et al. (2006). TRAIL receptor–mediated JNK activation and Bim phosphorylation critically regulate Fas-mediated liver damage and lethality. J. Clin. Invest. 116, 2493–2499. https://doi.org/10.1172/JCI27726.

Coste, A., Dubuquoy, L., Barnouin, R., Annicotte, J.-S., Magnier, B., Notti, M., Corazza, N., Antal, M.C., Metzger, D., Desreumaux, P., et al. (2007). LRH-1-mediated glucocorticoid synthesis in enterocytes protects against inflammatory bowel disease. Proc. Natl. Acad. Sci. 104, 13098–13103. https://doi.org/10.1073/pnas.0702440104.

Dubé, C., Bergeron, F., Vaillant, M.J., Robert, N.M., Brousseau, C., and Tremblay, J.J. (2009). The nuclear receptors SF1 and LRH1 are expressed in endometrial cancer cells and regulate steroidogenic gene transcription by cooperating with AP-1 factors. Cancer Lett. 275, 127–138. https://doi.org/10.1016/j.canlet.2008.10.008.

Duma, D., Collins, J.B., Chou, J.W., and Cidlowski, J.A. (2010). Sexually dimorphic actions of glucocorticoids provide a link to inflammatory diseases with gender differences in prevalence. Sci. Signal. 3, 1–12. https://doi.org/10.1126/scisignal.2001077.

Feng, R., Liebe, R., and Weng, H.-L. (2023). Transcription networks in liver development and acute liver failure. Liver Res. 7, 47–55. https://doi.org/10.1016/j.livres.2022.11.010.

Fernandez-Marcos, P.J., Auwerx, J., and Schoonjans, K. (2011). Emerging actions of the nuclear receptor LRH-1 in the gut. Biochim. Biophys. Acta - Mol. Basis Dis. 1812, 947–955. https://doi.org/10.1016/j.bbadis.2010.12.010.

Gionet, N., Jansson, D., Mader, S., and Pratt, M.A.C. (2009). NF-κB and estrogen receptor α interactions: Differential function in estrogen receptor-negative and -positive hormone-independent breast cancer cells. J. Cell. Biochem. 107, 448–459. https://doi.org/10.1002/jcb.22141.

Hamesch, k., Borkham-Kamphorst, E., Strnad, P., and Weiskirchen, R. (2015). Lipopolysaccharide-induced inflammatory liver injury in mice. Lab. Anim. 49, 37–46. https://doi.org/10.1177/0023677215570087.

Hattori, T., Iizuka, K., Horikawa, Y., and Takeda, J. (2014). LRH-1 heterozygous knockout mice are prone to mild obesity. Endocr. J. 61, 471–480. https://doi.org/10.1507/endocrj.EJ14-0017.

Heymann, F., Hamesch, K., Weiskirchen, R., and Tacke, F. (2015). The concanavalin A model of acute hepatitis in mice. Lab. Anim. 49, 12–20. https://doi.org/10.1177/0023677215572841.

Holbrook, J., Lara-Reyna, S., Jarosz-Griffiths, H., and McDermott, M.F. (2019). Tumour necrosis factor signalling in health and disease. F1000Research 8, 111. https://doi.org/10.12688/f1000research.17023.1.

Huang, S., Lee, C., and Chung, B. (2014). Tumor Necrosis Factor Suppresses NR5A2 Activity and Intestinal Glucocorticoid Synthesis to Sustain Chronic Colitis. Sci. Signal. 7, ra20. https://doi.org/10.1126/scisignal.2004786.

Hubler, M.J., and Kennedy, A.J. (2016). Role of lipids in the metabolism and activation of immune cells. J. Nutr. Biochem. 34, 1–7. https://doi.org/10.1016/j.jnutbio.2015.11.002.

Ingelfinger, F., Krishnarajah, S., Kramer, M., Utz, S.G., Galli, E., Lutz, M., Zwicky, P., Akarca, A.U., Jurado, N.P., Ulutekin, C., et al. (2021). Single-cell profiling of myasthenia gravis identifies a pathogenic T cell signature. Acta Neuropathol. 141, 901–915. https://doi.org/10.1007/s00401-021-02299-y.

Joo, M.S., Koo, J.H., Kim, T.H., Kim, Y.S., and Kim, S.G. (2019). LRH1-driven transcription factor circuitry for hepatocyte identity: Super-enhancer cistromic analysis. EBioMedicine 40, 488–503. https://doi.org/10.1016/j.ebiom.2018.12.056.

Jost, P.J., Grabow, S., Gray, D., McKenzie, M.D., Nachbur, U., Huang, D.C.S., Bouillet, P., Thomas, H.E., Borner, C., Silke, J., et al. (2009). XIAP discriminates between type I and type II FAS-induced apoptosis. Nature 460, 1035–1039. https://doi.org/10.1038/nature08229.

Karin, M., and Lin, A. (2002). NF-kappaB at the crossroads of life and death. Nat. Immunol. 3, 221–227. https://doi.org/10.1038/ni0302-221.

Kaufmann, T., Jost, P.J., Pellegrini, M., Puthalakath, H., Gugasyan, R., Gerondakis, S., Cretney, E., Smyth, M.J., Silke, J., Hakem, R., et al. (2009). Fatal Hepatitis Mediated by Tumor Necrosis Factor TNFα Requires Caspase-8 and Involves the BH3-Only Proteins Bid and Bim. Immunity 30, 56–66. https://doi.org/10.1016/j.immuni.2008.10.017.

Kaufmann, T., Strasser, A., and Jost, P.J. (2012). Fas death receptor signalling: roles of Bid and XIAP. Cell Death Differ. 19, 42–50. https://doi.org/10.1038/cdd.2011.121.

Kietzmann, T. (2019). Liver Zonation in Health and Disease: Hypoxia and Hypoxia-Inducible Transcription Factors as Concert Masters. Int. J. Mol. Sci. 20, 2347. https://doi.org/10.3390/ijms20092347.

Klatt, K.C., Petviashvili, E.J., and Moore, D.D. (2023). LRH-1 induces hepatoprotective nonessential amino acids in response to acute liver injury. J. Clin. Invest. 133, 5–8. https://doi.org/10.1172/JCI168805.

Ksontini, R., Colagiovanni, D.B., Josephs, M.D., Edwards, C.K., Tannahill, C.L., Solorzano, C.C., Norman, J., Denham, W., Clare-Salzler, M., MacKay, S.L.D., et al. (1998). Disparate Roles for TNF-α and Fas Ligand in Concanavalin A-Induced Hepatitis. J. Immunol. 160, 4082–4089. https://doi.org/10.4049/jimmunol.160.8.4082.

Lai, C.-F., Flach, K.D., Alexi, X., Fox, S.P., Ottaviani, S., Thiruchelvam, P.T.R., Kyle, F.J., Thomas, R.S., Launchbury, R., Hua, H., et al. (2013). Co-regulated gene expression by oestrogen receptor α and liver receptor homolog-1 is a feature of the oestrogen response in breast cancer cells. Nucleic Acids Res. 41, 10228–10240. https://doi.org/10.1093/nar/gkt827.

Lazarus, K.A., Wijayakumara, D., Chand, A.L., Simpson, E.R., and Clyne, C.D. (2012). Therapeutic potential of Liver Receptor Homolog-1 modulators. J. Steroid Biochem. Mol. Biol. 130, 138–146. https://doi.org/10.1016/j.jsbmb.2011.12.017.

Lefèvre, L., Authier, H., Stein, S., Majorel, C., Couderc, B., Dardenne, C., Eddine, M.A., Meunier, E., Bernad, J., Valentin, A., et al. (2015). LRH-1 mediates anti-inflammatory and antifungal phenotype of IL-13-activated macrophages through the PPARγ ligand synthesis. Nat. Commun. 6. https://doi.org/10.1038/ncomms7801.

Leist, M., Gantner, F., Bohlinger, I., Germann, P.G., Tiegs, G., and Wendel, A. (1994). Murine hepatocyte apoptosis induced in vitro and in vivo by TNF-alpha requires transcriptional arrest. J. Immunol. 153, 1778–1788. https://doi.org/10.4049/jimmunol.153.4.1778.

Liu, Z., Gu, Y., Shin, A., Zhang, S., and Ginhoux, F. (2020). Analysis of Myeloid Cells in Mouse Tissues with Flow Cytometry. STAR Protoc. 1, 100029. https://doi.org/10.1016/j.xpro.2020.100029.

Loft, A., Alfaro, A.J., Schmidt, S.F., Pedersen, F.B., Terkelsen, M.K., Puglia, M., Chow, K.K., Feuchtinger, A., Troullinaki, M., Maida, A., et al. (2021). Liver-fibrosis-activated transcriptional networks govern hepatocyte reprogramming and intra-hepatic communication. Cell Metab. 33, 1685–1700.e9. https://doi.org/10.1016/j.cmet.2021.06.005.

Loft, A., Schmidt, S.F., Caratti, G., Stifel, U., Havelund, J., Sekar, R., Kwon, Y., Sulaj, A., Chow, K.K., Alfaro, A.J., et al. (2022). A macrophage-hepatocyte glucocorticoid receptor axis coordinates fasting ketogenesis. Cell Metab. 34, 473–486.e9. https://doi.org/10.1016/j.cmet.2022.01.004.

Lu, T.T., Makishima, M., Repa, J.J., Schoonjans, K., Kerr, T.A., Auwerx, J., and Mangelsdorf, D.J. (2000). Molecular Basis for Feedback Regulation of Bile Acid Synthesis by Nuclear Receptors. Mol. Cell 6, 507–515. https://doi.org/10.1016/S1097-2765(00)00050-2.

Luciano, F., Jacquel, A., Colosetti, P., Herrant, M., Cagnol, S., Pages, G., and Auberger, P. (2003). Phosphorylation of Bim-EL by Erk1/2 on serine 69 promotes its degradation via the proteasome pathway and regulates its proapoptotic function. Oncogene 22, 6785–6793. https://doi.org/10.1038/sj.onc.1206792.

Martin Vázquez, E., Cobo-Vuilleumier, N., Araujo Legido, R., Marín-Cañas, S., Nola, E., Dorronsoro, A., López Bermudo, L., Crespo, A., Romero-Zerbo, S.Y., García-Fernández, M., et al. (2022). NR5A2/LRH-1 regulates the PTGS2-PGE2-PTGER1 pathway contributing to pancreatic islet survival and function. IScience 25. https://doi.org/10.1016/j.isci.2022.104345.

Mataki, C., Magnier, B.C., Houten, S.M., Annicotte, J.-S., Argmann, C., Thomas, C., Overmars, H., Kulik, W., Metzger, D., Auwerx, J., et al. (2007). Compromised Intestinal Lipid Absorption in Mice with a Liver-Specific Deficiency of Liver Receptor Homolog 1. Mol. Cell. Biol. 27, 8330–8339. https://doi.org/10.1128/MCB.00852-07.

Matsukuma, K.E., Wang, L., Bennett, M.K., and Osborne, T.F. (2007). A Key Role for Orphan Nuclear Receptor Liver Receptor Homologue-1 in Activation of Fatty Acid Synthase Promoter by Liver X Receptor. J. Biol. Chem. 282, 20164–20171. https://doi.org/10.1074/jbc.M702895200.

Mehlem, A., Hagberg, C.E., Muhl, L., Eriksson, U., and Falkevall, A. (2013). Imaging of neutral lipids by oil red O for analyzing the metabolic status in health and disease. Nat. Protoc. 8, 1149–1154. https://doi.org/10.1038/nprot.2013.055.

Meinsohn, M.-C., Smith, O.E., Bertolin, K., and Murphy, B.D. (2019). The Orphan Nuclear Receptors Steroidogenic Factor-1 and Liver Receptor Homolog-1: Structure, Regulation, and Essential Roles in Mammalian Reproduction. Physiol. Rev. 99, 1249–1279. https://doi.org/10.1152/physrev.00019.2018.

Michalek, S., Goj, T., Plazzo, A.P., Marovca, B., Bornhauser, B., and Brunner, T. (2022). LRH-1/NR5A2 interacts with the glucocorticoid receptor to regulate glucocorticoid resistance. EMBO Rep. 23, e54195. https://doi.org/10.15252/embr.202154195.

Miranda, D.A., Krause, W.C., Cazenave-Gassiot, A., Suzawa, M., Escusa, H., Foo, J.C., Shihadih, D.S., Stahl, A., Fitch, M., Nyangau, E., et al. (2018). LRH-1 regulates hepatic lipid homeostasis and maintains arachidonoyl phospholipid pools critical for phospholipid diversity. JCI Insight 3. https://doi.org/10.1172/jci.insight.96151.

Mohamad, N.-V., Wong, S.K., Wan Hasan, W.N., Jolly, J.J., Nur-Farhana, M.F., Ima-Nirwana, S., and Chin, K.-Y. (2019). The relationship between circulating testosterone and inflammatory cytokines in men. Aging Male 22, 129–140. https://doi.org/10.1080/13685538.2018.1482487.

Mossanen, J.C., and Tacke, F. (2015). Acetaminophen-induced acute liver injury in mice. Lab. Anim. 49, 30–36. https://doi.org/10.1177/0023677215570992.

Mueller, M., Cima, I., Noti, M., Fuhrer, A., Jakob, S., Dubuquoy, L., Schoonjans, K., and Brunner, T. (2006). The nuclear receptor LRH-1 critically regulates extra-adrenal glucocorticoid synthesis in the intestine. J. Exp. Med. 203, 2057–2062. https://doi.org/10.1084/jem.20060357.

Nadolny, C., and Dong, X. (2015). Liver receptor homolog-1 (LRH-1): a potential therapeutic target for cancer. Cancer Biol. Ther. 16, 997–1004. https://doi.org/10.1080/15384047.2015.1045693.

Nowicka, M., Krieg, C., Crowell, H.L., Weber, L.M., Hartmann, F.J., Guglietta, S., Becher, B., Levesque, M.P., and Robinson, M.D. (2019). CyTOF workflow: differential discovery in high-throughput high-dimensional cytometry datasets. F1000Research 6, 748. https://doi.org/10.12688/f1000research.11622.3.

Oosterveer, M.H., and Schoonjans, K. (2014). Hepatic glucose sensing and integrative pathways in the liver. Cell. Mol. Life Sci. 71, 1453–1467. https://doi.org/10.1007/s00018-013-1505-z.

Oosterveer, M.H., Mataki, C., Yamamoto, H., Harach, T., Moullan, N., van Dijk, T.H., Ayuso, E., Bosch, F., Postic, C., Groen, A.K., et al. (2012). LRH-1–dependent glucose sensing determines intermediary metabolism in liver. J. Clin. Invest. 122, 2817–2826. https://doi.org/10.1172/JCI62368.

Ortlund, E.A., Lee, Y., Solomon, I.H., Hager, J.M., Safi, R., Choi, Y., Guan, Z., Tripathy, A., Raetz, C.R.H., McDonnell, D.P., et al. (2005). Modulation of human nuclear receptor LRH-1 activity by phospholipids and SHP. Nat. Struct. Mol. Biol. 12, 357–363. https://doi.org/10.1038/nsmb910.

Paré, J.-F., Malenfant, D., Courtemanche, C., Jacob-Wagner, M., Roy, S., Allard, D., and Bélanger, L. (2004). The Fetoprotein Transcription Factor (FTF) Gene Is Essential to Embryogenesis and Cholesterol Homeostasis and Is Regulated by a DR4 Element. J. Biol. Chem. 279, 21206–21216. https://doi.org/10.1074/jbc.M401523200.

Phan, T.S., Schink, L., Mann, J., Merk, V.M., Zwicky, P., Mundt, S., Simon, D., Kulms, D., Abraham, S., Legler, D.F., et al. (2021). Keratinocytes control skin immune homeostasis through de novo– synthesized glucocorticoids. Sci. Adv. 7, 1–20. https://doi.org/10.1126/sciadv.abe0337.

Potter, D.S., and Letai, A. (2016). To Prime, or Not to Prime: That Is the Question. Cold Spring Harb. Symp. Quant. Biol. 81, 131–140. https://doi.org/10.1101/sqb.2016.81.030841.

Ramachandran, A., and Jaeschke, H. (2019a). Acetaminophen hepatotoxicity: A mitochondrial perspective. In Advances in Pharmacology, (Elsevier Inc.), pp. 195–219.

Ramachandran, A., and Jaeschke, H. (2019b). Acetaminophen Hepatotoxicity. Semin. Liver Dis. 39, 221–234. https://doi.org/10.1055/s-0039-1679919.

Schoonjans, K., Annicotte, J., Huby, T., Botrugno, O.A., Fayard, E., Ueda, Y., Chapman, J., and Auwerx, J. (2002). Liver receptor homolog 1 controls the expression of the scavenger receptor class B type I. EMBO Rep. 3, 1181–1187. https://doi.org/10.1093/embo-reports/kvf238.

Schoonjans, K., Dubuquoy, L., Mebis, J., Fayard, E., Wendling, O., Haby, C., Geboes, K., and Auwerx, J. (2005). Liver receptor homolog 1 contributes to intestinal tumor formation through effects on cell cycle and inflammation. Proc. Natl. Acad. Sci. 102, 2058–2062. https://doi.org/10.1073/pnas.0409756102.

Schwaderer, J., Gaiser, A.-K., Phan, T.S., Delgado, M.E., and Brunner, T. (2017). Liver receptor homolog-1 (NR5a2) regulates CD95/Fas ligand transcription and associated T-cell effector functions. Cell Death Dis. 8, e2745–e2745. https://doi.org/10.1038/cddis.2017.173.

Schwaderer, J., Phan, T.S., Glöckner, A., Delp, J., Leist, M., Brunner, T., and Delgado, M.E. (2020). Pharmacological LRH-1/Nr5a2 inhibition limits pro-inflammatory cytokine production in macrophages and associated experimental hepatitis. Cell Death Dis. 11, 154. https://doi.org/10.1038/s41419-020-2348-9.

Seitz, C., Huang, J., Geiselhöringer, A.-L., Galbani-Bianchi, P., Michalek, S., Phan, T.S., Reinhold, C., Dietrich, L., Schmidt, C., Corazza, N., et al. (2019). The orphan nuclear receptor LRH-1/NR5a2 critically regulates T cell functions. Sci. Adv. 5, eaav9732. https://doi.org/10.1126/sciadv.aav9732.

Semjonous, N.M., Sherlock, M., Jeyasuria, P., Parker, K.L., Walker, E.A., Stewart, P.M., and Lavery, G.G. (2011). Hexose-6-Phosphate Dehydrogenase Contributes to Skeletal Muscle Homeostasis Independent of 11β-Hydroxysteroid Dehydrogenase Type 1. Endocrinology 152, 93–102. https://doi.org/10.1210/en.2010-0957.

Sia, D., Villanueva, A., Friedman, S.L., and Llovet, J.M. (2017). Liver Cancer Cell of Origin, Molecular Class, and Effects on Patient Prognosis. Gastroenterology 152, 745–761. https://doi.org/10.1053/j.gastro.2016.11.048.

Sirianni, R., Seely, J., Attia, G., Stocco, D., Carr, B., Pezzi, V., and Rainey, W. (2002). Liver receptor homologue-1 is expressed in human steroidogenic tissues and activates transcription of genes encoding steroidogenic enzymes. J. Endocrinol. 174, R13–R17. https://doi.org/10.1677/joe.0.174r013.

Stein, S., Lemos, V., Xu, P., Demagny, H., Wang, X., Ryu, D., Jimenez, V., Bosch, F., Lüscher, T.F., Oosterveer, M.H., et al. (2017). Impaired SUMOylation of nuclear receptor LRH-1 promotes nonalcoholic fatty liver disease. J. Clin. Invest. 127, 583–592. https://doi.org/10.1172/JCI85499.

Straub, R.H. (2007). The Complex Role of Estrogens in Inflammation. Endocr. Rev. 28, 521–574. https://doi.org/10.1210/er.2007-0001.

Sun, Y., Demagny, H., and Schoonjans, K. (2021). Emerging functions of the nuclear receptor LRH-1 in liver physiology and pathology. Biochim. Biophys. Acta - Mol. Basis Dis. 1867, 166145. https://doi.org/10.1016/j.bbadis.2021.166145.

Sun, Y., Demagny, H., Faure, A., Pontanari, F., Jalil, A., Bresciani, N., Yildiz, E., Korbelius, M., Perino, A., and Schoonjans, K. (2023). Asparagine protects pericentral hepatocytes during acute liver injury. J. Clin. Invest. https://doi.org/10.1172/JCI163508.

Thiruchelvam, P.T.R., Lai, C.-F., Hua, H., Thomas, R.S., Hurtado, A., Hudson, W., Bayly, A.R., Kyle, F.J., Periyasamy, M., Photiou, A., et al. (2011). The liver receptor homolog-1 regulates estrogen receptor expression in breast cancer cells. Breast Cancer Res. Treat. 127, 385–396. https://doi.org/10.1007/s10549-010-0994-9.

Venteclef, N., Smith, J.C., Goodwin, B., and Delerive, P. (2006). Liver Receptor Homolog 1 Is a Negative Regulator of the Hepatic Acute-Phase Response. Mol. Cell. Biol. 26, 6799–6807. https://doi.org/10.1128/MCB.00579-06.

Vince, J.E., Wong, W.W.L., Khan, N., Feltham, R., Chau, D., Ahmed, A.U., Benetatos, C.A., Chunduru, S.K., Condon, S.M., McKinlay, M., et al. (2007). IAP Antagonists Target cIAP1 to Induce TNFα-Dependent Apoptosis. Cell 131, 682–693. https://doi.org/10.1016/j.cell.2007.10.037.

Volle, D.H., Duggavathi, R., Magnier, B.C., Houten, S.M., Cummins, C.L., Lobaccaro, J.-M.A., Verhoeven, G., Schoonjans, K., and Auwerx, J. (2007). The small heterodimer partner is a gonadal gatekeeper of sexual maturation in male mice. Genes Dev. 21, 303–315. https://doi.org/10.1101/gad.409307.

Walesky, C.M., Kolb, K.E., Winston, C.L., Henderson, J., Kruft, B., Fleming, I., Ko, S., Monga, S.P., Mueller, F., Apte, U., et al. (2020). Functional compensation precedes recovery of tissue mass following acute liver injury. Nat. Commun. 11, 5785. https://doi.org/10.1038/s41467-020-19558-3.

Wang, J., Zhang, L., Shi, Q., Yang, B., He, Q., Wang, J., and Weng, Q. (2022). Targeting innate immune responses to attenuate acetaminophen-induced hepatotoxicity. Biochem. Pharmacol. 202, 115142. https://doi.org/10.1016/j.bcp.2022.115142.

Wang, S., Lan, F., Huang, L., Dong, L., Zhu, Z., Li, Z., Xie, Y., and Fu, J. (2005). Suppression of hLRH-1 mediated by a DNA vector-based RNA interference results in cell cycle arrest and induction of apoptosis in hepatocellular carcinoma cell BEL-7402. Biochem. Biophys. Res. Commun. 333, 917– 924. https://doi.org/10.1016/j.bbrc.2005.05.186.

Xiao, L., Wang, Y., Xu, K., Hu, H., Xu, Z., Wu, D., Wang, Z., You, W., Ng, C.-F., Yu, S., et al. (2018). Nuclear Receptor LRH-1 Functions to Promote Castration-Resistant Growth of Prostate Cancer via Its Promotion of Intratumoral Androgen Biosynthesis. Cancer Res. 78, 2205–2218. https://doi.org/10.1158/0008-5472.CAN-17-2341.

Xu, P., Oosterveer, M.H., Stein, S., Demagny, H., Ryu, D., Moullan, N., Wang, X., Can, E., Zamboni, N., Comment, A., et al. (2016). LRH-1-dependent programming of mitochondrial glutamine processing drives liver cancer. Genes Dev. 30, 1255–1260. https://doi.org/10.1101/gad.277483.116.

Zhou, J., Suzuki, T., Kovacic, A., Saito, R., Miki, Y., Ishida, T., Moriya, T., Simpson, E.R., Sasano, H., and Clyne, C.D. (2005). Interactions between Prostaglandin E2, Liver Receptor Homologue-1, and Aromatase in Breast Cancer. Cancer Res. 65, 657–663. https://doi.org/10.1158/0008-5472.657.65.2.

Zhou, J., Wang, Y., Wu, D., Wang, S., Chen, Z., Xiang, S., and Chan, F.L. (2021). Orphan nuclear receptors as regulators of intratumoral androgen biosynthesis in castration-resistant prostate cancer. Oncogene 40, 2625–2634. https://doi.org/10.1038/s41388-021-01737-1.

